# A genome-wide in vivo CRISPR screen identifies neuroprotective strategies in the mouse and human retina

**DOI:** 10.1101/2025.03.22.644712

**Authors:** Ning Shen, Michael J. Fitzpatrick, Ellen G. Harding, Sohini Rebba, Philip A. Ruzycki, Daniel Kerschensteiner

## Abstract

Retinitis pigmentosa (RP) is a genetically diverse blinding disorder lacking broadly effective therapies. We performed a genome-wide in vivo CRISPR knockout screen in mice carrying the *P23H* rhodopsin mutation (the most common cause of autosomal dominant RP in the United States) to systematically identify neuroprotective genes. We discovered multiple knockouts that accelerated rod photoreceptor loss, validated top candidates, and showed that overexpressing two genes—*UFD1* and *UXT*—preserved rods and cones, maintained retinal function, and improved visual behaviors. To accelerate translation, we developed a human *P23H* RP model in adult retinal explants, recreating key disease features. *UFD1* and *UXT* augmentation prevented photoreceptor loss in human *P23H* retinas. Our findings establish a pipeline for systematic identification and translational testing of neuroprotective genes in mouse and human RP models, provide a novel set of validated candidate genes, and underscore the therapeutic promise of *UFD1* and *UXT* as mutation-agnostic strategies to preserve vision.

## Introduction

Inherited retinal degenerations (IRDs), characterized by the progressive death and dysfunction of photoreceptors, cause visual impairment, including blindness, in young adults ^1, 2^. IRDs are genetically the most heterogenous group of heritable disorders in humans. Mutations in more than 300 genes are known to cause IRDs, and multiple disease-causing mutations have been identified in many genes ^3^. Thus, more than 150 mutations in rhodopsin cause RP, an IRD in which rods die first, followed by cones ^4^. The mutations underlying approximately one-third of IRDs remain unknown ^5, 6^. While most individual mutations are rare, collectively, IRDs are common, affecting millions of people worldwide (prevalence: ∼1 in 2,000) ^7, 8^. There are currently no effective treatments for most IRDs ^2, 9, 10^.

Despite their genetic diversity, IRDs converge on few pathogenic pathways, including protein misfolding and endoplasmic reticulum (ER) stress ^11^—a mechanism also implicated in neurodegenerative disease outside the eye, including Alzheimer’s disease ^12–14^, Huntington’s disease ^15, 16^, and amyotrophic lateral sclerosis ^14, 17^. Because IRD-causing mutations are individually rare but collectively impose a substantial disease burden, and because diverse mutations share common pathogenic pathways, IRDs are ideal candidates for mutation-agnostic neuroprotective therapies. To date, however, the search for neuroprotection—in the retina and the brain—has been hypothesis-driven and narrow in scope ^18, 19^. The arrival of CRISPR gene editing raises the possibility of comprehensive and unbiased screens to identify new therapeutic targets. Indeed, genome-wide CRISPR screens have identified novel candidates for treating cancer ^20–22^, autoimmune disorders ^23, 24^, and infectious diseases ^25–27^. In vivo CRISPR screens in the nervous system are rare ^28, 29^ and, this powerful approach has not yet been applied to the search for neuroprotection.

Here, we conduct a genome-wide in vivo CRISPR screen in rod photoreceptors to evaluate the neuroprotective potential of every protein-coding gene in the genome. We performed this screen in mice carrying the rhodopsin *P23H* mutation, the most common cause of autosomal dominant RP in the United States (accounting for 15–18% of cases), which leads to photoreceptor degeneration through rhodopsin misfolding and ER stress ^4, 30–33^. Our genome-wide screen identified a set of neuroprotective candidate genes with potential for therapeutic applications in the eye and beyond. A secondary screen confirmed top hits, and we developed gene therapy vectors for two candidates. These vectors effectively preserved retinal structure, function, and visual behaviors in *P23H* mice.

A major bottleneck in translation is the transition from animal models to humans. Human tissue systems can accelerate translation and help explore neurodegenerative disease mechanisms and neuroprotective therapies ^34–36^. Stem-cell-derived organoids, first developed in the retina^37, 38^, can incorporate patient mutations, but yield immature neurons and imprecise circuits, limiting their utility for modeling adult-onset diseases and synaptic deficits, which contribute to the progression of degenerative diseases ^39–42^. To overcome these limitations, we used viral engineering in retinal explant cultures from adult organ donors to develop an experimentally accessible human *P23H* model. We used this model to evaluate the translational potential of the gene therapies we developed in mice.

## Results

### Genome-wide in vivo CRISPR screen in *P23H* mice

To optimize our genome-wide CRISPR screen for identifying neuroprotective genes, we first characterized the time course of photoreceptor degeneration in heterozygous *P23H* mice expressing Cas9 in rods (*P23H Nrl-Cas9* mice; Fig. 1a,b). Consistent with prior observations ^33, 43^, the outer nuclear layer (ONL) thickness in these mice was comparable to wild-type (WT) controls during development. However, by one month of age, the ONL thickness had decreased to ∼74% of WT, and it continued to decline to ∼38% of WT by four months (Fig. 1a,b). We reasoned that a CRISPR-mediated knockout of neuroprotective genes would accelerate degeneration, causing depletion of rod photoreceptors carrying the corresponding single-guide RNAs (sgRNAs). To enhance the detection of sgRNA depletion (negative selection) and capture the most important neuroprotective factors, we chose a one-month endpoint—when degeneration is evident but not so advanced as to surface less critical genes and introduce confounds from bystander death.

**Fig. 1.**
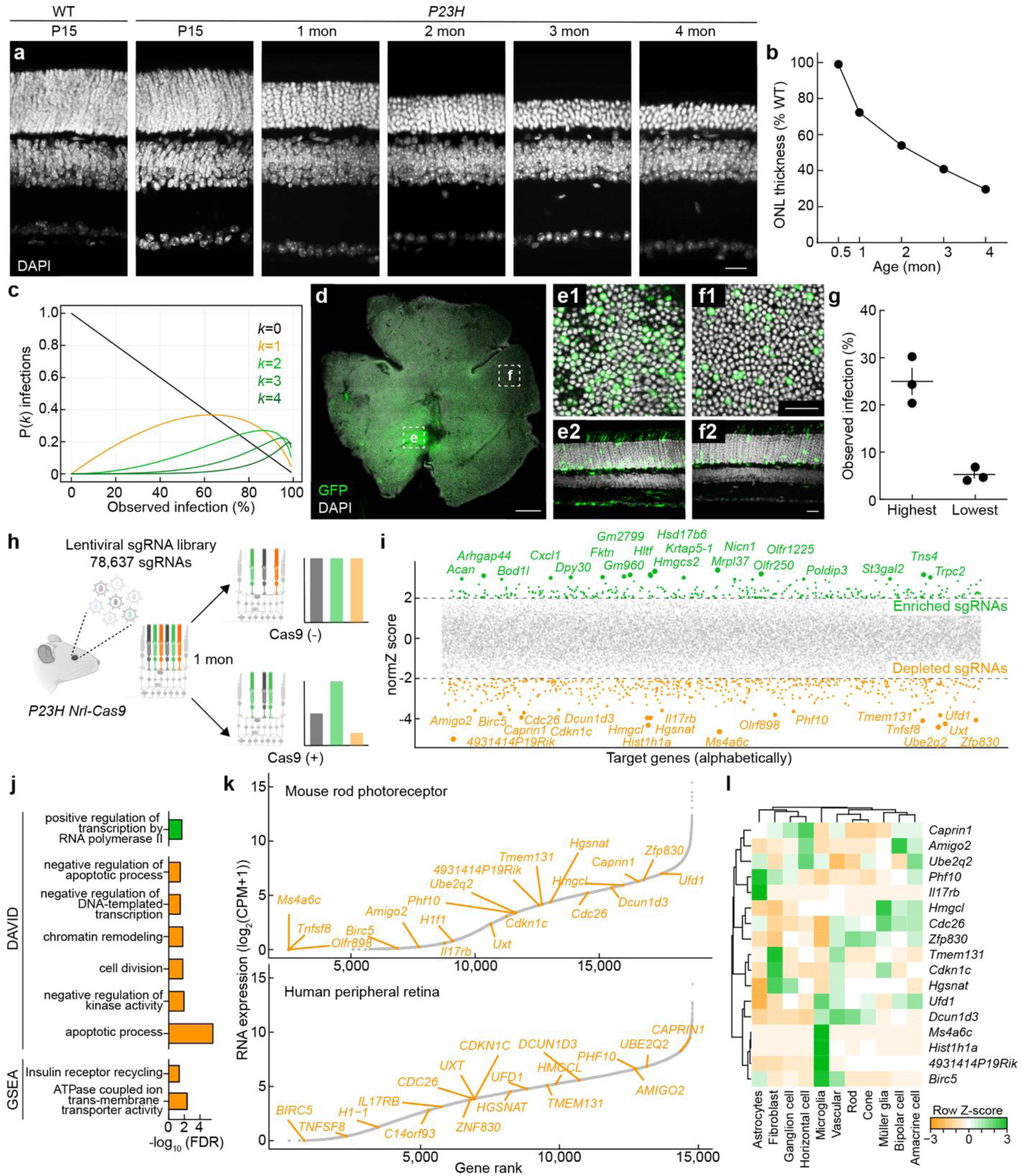
Genome-wide in vivo CRISPR screen in *P23H* mice. **a,b,** Time course of photoreceptor degeneration in *P23H* mice. (**a**) Representative vertical sections WT and heterozygous *P23H* retinas (DAPI stained) at postnatal day 15 (P15) and 1, 2, 3, and 4 months. Scale bar, 20 µm. (**b**) ONL thickness in *P23H* retinas normalized to WT littermates at each age. **c–g,** LV delivery in the mouse retina. (**c**) Estimates of the distributions of LV multiplicities of infection (i.e., fraction of cells infected by 0, 1, 2, etc. viruses) based on the observed labeling rate (see Methods). (**d**) Whole-mount image of an *LV–EF1α–GFP*-infected retina, with regions of highest (**e**) and lowest (**f**) transduction. (**e1**,**e2**,**f1**,**f2**) Enlarged whole-mount (**e1**,**f1**) and vertical-section (**e2**,**f2**) views of the areas in (**e,f**). Scale bars, 500 µm (**d**), 20 µm (**e1**,**e2**,**f1**,**f2**). (**g**) Quantification of labeling rates in the regions in high- and low-density regions (n = 3 retinas). **h–j**, Genome-wide in vivo CRISPR screen logic and outcomes. (**h**) Schematic of the screen. The Brie library ^44^, containing four sgRNAs per gene for the protein-coding mouse genome, was packaged into LVs and delivered subretinally to newborn *P23H* mice with rod-specific Cas9 expression. After one month, genomic DNA was extracted and sgRNA distributions sequenced. sgRNAs targeting neuroprotective genes become depleted (rod death is accelerated), while those targeting disease-promoting genes become enriched. (**i**) The comparison between *P23H* mice with (Cas9(+)) and without (Cas9(–)) rod-specific Cas9 yields normZ scores ^45^ for each gene (negative = depletion, positive = enrichment). Genes with |normZ| > 2 are color-coded (orange = depleted, green = enriched), with the top 20 annotated. **(j)** Gene Ontology (GO) and pathway analyses using DAVID (|normZ| > 2) and GSEA (ranked list of all genes), revealing enrichment of pathways related to apoptosis, chromatin regulation, and vacuolar ATPase-mediated lysosomal function. **k,l,** Expression profiles of the top 20 depleted genes in bulk RNA sequencing data from sorted mouse rods and the rod-rich peripheral human retina (**k**), and distribution of RNA expression across mouse retinal cell types (**l**). Data were obtained from SRR3662498, SRR3662513, and SRR3662514 for sorted mouse rods ^56^, SRR5591599 for peripheral human retina ^57^, and GSE63473 for mouse single-cell data ^93^.

Next, we optimized lentiviral (LV) delivery to infect as many rods as possible while minimizing the fraction of cells receiving more than one viral particle (and, thus, more than one sgRNA). We performed subretinal injections of a control LV expressing GFP (210 nL at postnatal day 1 [P1]) at a similar titer (1 × 10^9^ TU/mL) to our CRISPR library (1.26 × 10^9^ TU/mL) ^44^ and observed broad rod-dominant infection by one month (P30). As illustrated in Fig. 1c, the probability that a given cell is infected by *k* viruses can be deduced from the observed labeling fraction—i.e., the fraction of cells infected by at least one virus—assuming a Poisson process (see STAR Methods). We observed ∼25% labeling near the injection site and ∼5% in more distant regions (Fig. 1d–g). Assuming an even distribution of infection rates between high- and low-density regions, we estimate ∼900,000 rods per retina were infected by at least one LV, with ∼95,000 rods receiving more than one LV.

We verified that Cas9 expression in *Nrl-Cas9* mice is rod-specific and increases from P7 to P30 (Extended Data Fig. 1a,b). In parallel, LV-driven gene expression following P1 injections was undetectable at P7 but robust by P30 (Extended Data Fig. 1c–f). Consequently, CRISPR-mediated gene knockouts in our screen should occur primarily in differentiated rod photoreceptors of *P23H Nrl-Cas9* mice, with minimal impact on precursor cells. This approach therefore enables us to identify neuroprotective genes that act during early rod degeneration.

To comprehensively screen for neuroprotective targets, we used the Brie library covering 19,674 protein-coding genes in the mouse genome with four sgRNAs, plus 1,000 non-targeting controls (78,637 sgRNAs in total) ^44^. We obtained this library from Addgene (#73633), transformed it into *E. coli*, and amplified the resulting DNA for LV production. To ensure we had not lost complexity during this potential bottleneck, we amplified sgRNAs for sequencing from DNA taken before and after the bacterial expansion step. Sequencing of replicates from both libraries showed that the Gini coefficient was within an acceptable range (Extended Data Fig. 2a). We packaged the amplified library into LVs and delivered it subretinally (210 nL of 1.26 × 10^9^ TU/mL per eye) to newborn (P1) *P23H Nrl-Cas9* mice as well as *P23H* controls lacking *Nrl-Cas9* (Fig. 1h). Retinas were collected at one month (P30) for genomic DNA isolation. Only samples passing quality-control thresholds for robust lentiviral integration (see Methods) were processed for library preparation. In total, 31 and 44 were selected from 89 *P23H Nrl-Cas9* and 141 control injected eyes, respectively.

Amplified sgRNAs were quantified by next-generation sequencing (>100 million reads per library), resulting in robust coverage of >99% of sgRNAs (Extended Data Fig. 2b). These data enabled us to identify both positive and negative effects of target gene knockout by comparing abundance in *P23H Nrl-Cas9* vs. *P23H* control retinas with the drugZ pipeline ^45^ (Supplementary Tables 1 and 2). Because we were restricted to four sgRNAs per gene ^44^ and faced inherent challenges of negative selection—namely, that dropout requires large sgRNA effects and has a limited dynamic range ^44, 46, 47^, coupled with variations among photoreceptors due to the stochastic nature of in vivo degeneration ^48, 49^—only five individual targets surpassed stringent false-discovery-rate (FDR) thresholds, consistent with previous results ^29, 50^. Nonetheless, high-ranking genes (under more relaxed FDR thresholds) in genome-wide in vivo CRISPR screens often provide valuable insights into disease mechanisms and identify therapeutic targets (Fig. 1i) ^51–55^. We therefore rank-ordered our data based on each gene’s Z-score and performed Gene Ontology (GO) analysis of genes with cumulative Z-scores >|2|, as well as Gene Set Enrichment Analysis (GSEA) on the entire dataset (Fig. 1j; Extended Data Fig. 2c–e). We observed both positively selected targets (Z-score >2), whose knockout appeared to enhance survival, and negatively selected targets (Z-score <–2), whose knockout depleted rod photoreceptors—an outcome more prevalent by design. Pathways implicated in the negatively selected group included apoptosis, chromatin regulation, gene expression, and vacuolar ATPase– mediated lysosome acidification (Fig. 1j; Extended Data Fig. 2c–e). Although these pathways each hold therapeutic promise, we focused our subsequent validation efforts on the 20 most strongly depleted genes identified in the primary screen.

Because our screen aimed to test gene knockouts specifically in rod photoreceptors, we verified that 17 of the top 20 depleted genes were expressed in purified adult mouse rods, and that 18 identifiable human orthologs are transcribed in the rod-rich peripheral retina (Fig. 1k) ^56, 57^. Single-cell RNA-seq further revealed that most of these genes are broadly expressed across retinal neuron types and glia, suggesting they participate in core cellular functions rather than rod-specific processes (Fig. 1l).

Thus, we demonstrate the feasibility of a genome-wide in vivo CRISPR screen in the degenerating retina. Our results reveal a set of candidate neuroprotective genes whose loss accelerates photoreceptor death, highlighting potential new therapeutic targets for IRDs.

### Validating CRISPR screen results and testing for essential genes

To verify the reliability of our genome-wide screen and confirm that the most depleted genes are indeed critical for rod survival in *P23H* mice, we conducted a secondary screen targeting 10 of the top 20 most depleted genes (selecting the odd-ranked genes). We designed constructs that Cre-dependently co-express two gene-specific sgRNAs (distinct from those used in the primary screen) and a fluorescent reporter (tdTomato). We then delivered these constructs to the rod photoreceptors of newborn *P23H Nrl-Cas9* mice via subretinal injections followed by in vivo electroporation (Fig. 2a,b) ^58, 59^.

**Fig. 2.**
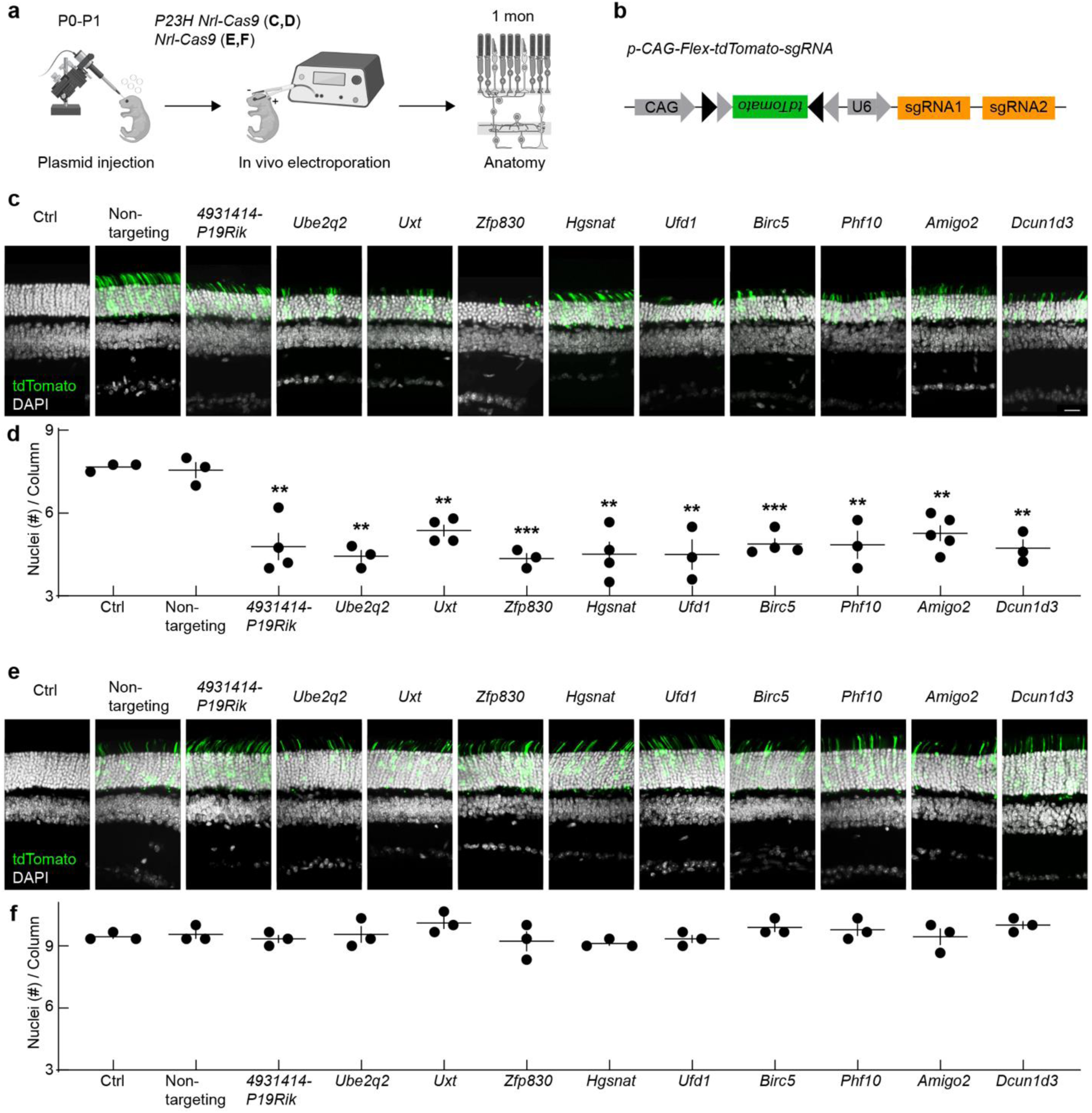
Validating CRISPR screen results and testing for essential genes. **a,b,** Schematic of the in vivo electroporation workflow (**a**) and dual–*sgRNA/tdTomato* construct (**b**) used to validate top-ranked CRISPR screen hits in newborn *P23H Nrl-Cas9* mice. Retinas were collected at one month. **c,** Representative confocal images of vertical retinal sections from *P23H Nrl-Cas9* mice without electroporation (Ctrl), or electroporated with plasmids encoding non-targeting sgRNAs (Non-targeting) or sgRNAs against ten odd-ranked top hits. tdTomato-positive cells (green) mark transduced photoreceptors; nuclei are stained with DAPI (white). Scale bar, 20 μm. **d,** Quantification of outer nuclear layer (ONL) nuclei per column from (**c**). **e,** As in (**c**), but in *Nrl-Cas9* mice lacking the *P23H* allele to assess whether these genes are essential in healthy rods. **f,** Quantification of ONL nuclei per column from (**e**). Data in (**d**) and (**f**) are plotted per retina (dots) with mean ± SEM (lines). One-way ANOVA compares each target to Non-targeting; **p < 0.01, ***p < 0.001, and no asterisk indicates p ≥ 0.05.

At one month of age, the control construct harboring non-targeting sgRNAs did not affect photoreceptor numbers (Fig. 2c,d). In contrast, each of the 10 gene-targeting constructs resulted in pronounced rod loss, reflected in a significant decrease in ONL thickness relative to non-targeting controls (Fig. 2c,d). To determine whether any of these genes were essential for rod survival under normal conditions, we repeated the secondary screen in *Nrl-Cas9* mice lacking the *P23H* mutation. None of the targeting constructs induced photoreceptor loss in these WT-like mice, indicating that the candidate genes are not required for rod viability in the absence of the *P23H* allele (Fig. 2e,f).

Collectively, these data validate our primary screen results and highlight that knocking out any of these 10 genes specifically exacerbates rod degeneration in the *P23H* context while sparing rods in the absence of the disease mutation. Our findings thus underscore the neuroprotective role of these genes and support the broader list of depleted genes from our primary screen as a promising resource for developing therapeutic strategies for IRDs.

### Neuroprotective gene therapies improve photoreceptor survival, morphology, and connectivity in ***P23H* mice**

Our primary and secondary CRISPR screens identified genes whose knockout accelerates photoreceptor degeneration in *P23H* mice. We hypothesized that enhancing the function of these genes could improve photoreceptor survival and function. To test this, we focused on two validated candidates, Ufd1 (Ubiquitin fusion degradation 1) and Uxt (Ubiquitously expressed transcript), and evaluated whether their overexpression could protect photoreceptors in *P23H* mice. The *P23H* mutation leads to misfolded rhodopsin and consequent ER stress ^32, 60, 61^. Ufd1 is a key component of the ER-associated degradation (ERAD) pathway, which removes misfolded proteins to mitigate ER stress ^62–64^. Uxt has been implicated in aggrephagy, which removes misfolded protein aggregates, and apoptosis regulation in neurons, including photoreceptors ^65–68^.

To increase the translational relevance of our neuroprotective approach, we cloned the human *UFD1* and *UXT* genes into adeno-associated viral (AAV) vectors (Fig. 3a). We then delivered them by subretinal injections to *P23H* mice and restricted expression to rods through Cre-mediated recombination (Fig. 3a,b). We confirmed robust viral infection, efficient recombination, and target gene expression by RT-PCR (Extended Data Fig. 3).

**Fig. 3.**
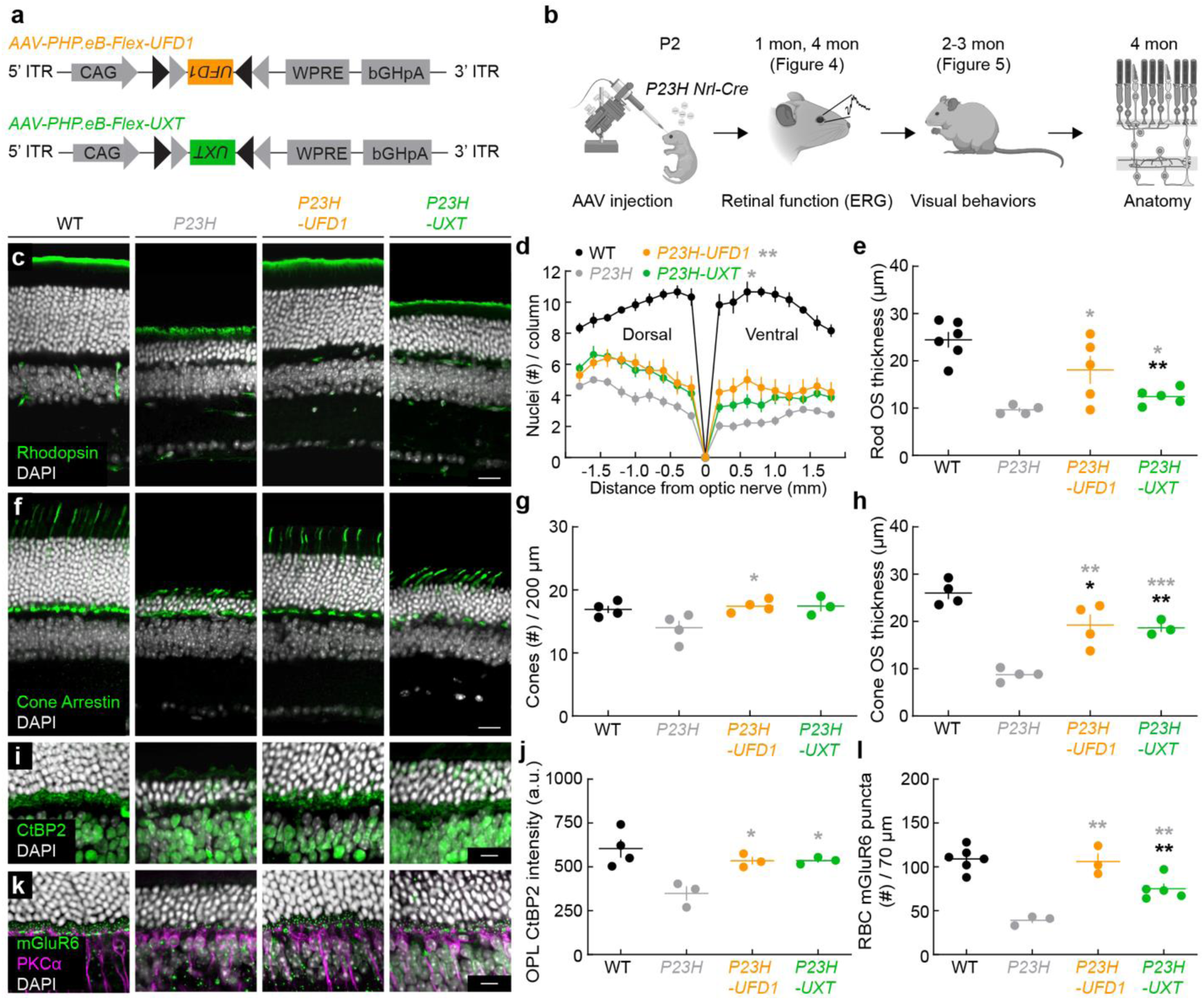
*UFD1* and *UXT* augmentation improve photoreceptor survival, morphology, and connectivity in *P23H* mice. **a,** Schematics of the AAV constructs (*AAV-PHP.eB-Flex*) used for rod-specific overexpression of *UFD1* or *UXT*. **b,** Experimental timeline for testing the effects of *UFD1* or *UXT* overexpression on retinal structure, function, and visual behaviors in *P23H* knockin mice. **c,** Representative confocal images of ventral retinal sections from wild-type (WT), untreated *P23H*, or *P23H* treated with *UFD1* (*P23H-UFD1*) or *UXT* (*P23H-UXT*), stained for rhodopsin (green) and DAPI (white). Scale bar, 20 µm. **d,** Number of ONL nuclei per column plotted as a function of distance from the optic nerve. Data are mean ± SEM; n = 7–12 retinas from 4–7 mice per group. Gray asterisks indicate significance compared to *P23H* (*p < 0.05, **p < 0.01) by Monte Carlo permutation testing. **e,** Quantification of rod outer segment (OS) thickness from (**c**). **f,** Representative images of ventral retinal sections from WT, *P23H*, *P23H-UFD1*, and *P23H-UXT* stained for cone arrestin (green) and DAPI (white). Scale bar, 20 µm. **g,h,** Summary data for cone density (**g**) and OS thickness (**h**). **i,** Representative images of ventral retinas stained for CtBP2 (green) and DAPI (white), labeling presynaptic ribbons. Scale bar, 10 µm. **j,** Quantification of CtBP2 immunofluorescence in the outer plexiform layer (OPL). **k,** Representative images stained for mGluR6 (green), PKCα (magenta), and DAPI (white). Scale bar, 10 µm. **l,** Quantification of mGluR6 puncta at rod bipolar cell dendrites. For (**e,g,h,j,l**), absence of an asterisk indicates p ≥ 0.05; black asterisks (*p < 0.05, **p < 0.01) indicate significance vs. WT, while gray asterisks (*p < 0.05, **p < 0.001, ***p < 0.001) indicate significance vs. *P23H* by one-way ANOVA.

We first evaluated whether *UFD1* or *UXT* overexpression ameliorated rod loss. Previous studies have shown that rod degeneration in *P23H* mice follows a retinal gradient (ventral > dorsal), and our data corroborated this pattern (Fig. 3d) ^33^. By four months, AAV-mediated overexpression of *UFD1* or *UXT* significantly improved rod survival across the retina (Fig. 3c,d). The remainder of Fig. 3 highlights the ventral retina; parallel findings for the dorsal retina are presented in Extended Data Fig. 4.

As rods degenerate, their outer segments shorten, contributing to visual deficits. Overexpression of *UFD1* or *UXT* prevented this shortening in *P23H* mice (Fig. 3c,e; Extended Data Fig. 4a,b). Because rods provide trophic and structural support to cones, rod degeneration ultimately compromises cone integrity ^69–71^. Although cone numbers in *P23H* mice remained stable at four months, their outer segments exhibited significant deterioration (Fig. 3f–h; Extended Data Fig. 4c–e). Overexpression of *UFD1* or *UXT* mitigated this cone outer segment loss, indicating a protective effect that extends beyond rods to preserve cone health as well.

Neurodegenerative diseases, including IRDs, frequently present early synapse loss that contributes to disease progression ^11, 41, 72, 73^. By four months, synapses between photoreceptors and second-order neurons were diminished in *P23H* mice, as evidenced by reductions in pre- and postsynaptic markers (Fig. 3i–3l; Extended Data Fig. 4f–i). Remarkably, *UFD1* and *UXT* overexpression preserved these synapses, preventing the loss of photoreceptor connections to downstream neurons.

Thus, our gene therapy approach targeting *UFD1* or *UXT* in *P23H* mice improved rod survival, prevented secondary cone degeneration, and maintained synaptic integrity.

### Neuroprotective gene therapies preserve light responses in *P23H* retinas

Having observed that *UFD1* or *UXT* overexpression improves rod survival and protects rod and cone outer segments (the sites of phototransduction) as well as synapses in *P23H* mice, we next asked whether these interventions improve retinal function. We evaluated both rod-driven (dark-adapted) and cone-driven (light-adapted) responses via in vivo electroretinography (ERG).

At one month, *P23H* mice exhibited significantly reduced rod-driven ERG amplitudes relative to WT controls (Fig. 4a–c) ^33, 43^. This decline affected both the a-wave (reflecting photoreceptor hyperpolarization; Fig. 4b) and the b-wave (reflecting ON bipolar cell depolarization; Fig. 4c). Overexpression of *UFD1* or *UXT* preserved these responses, resulting in a- and b-wave amplitudes that lay between those of WT and untreated *P23H* mice (Fig. 4a–c). Meanwhile, cone-driven ERG responses remained near WT levels in *P23H* mice at this early stage and were unaffected by *UFD1* or *UXT* gene therapy (Fig. 4d,e).

**Fig. 4.**
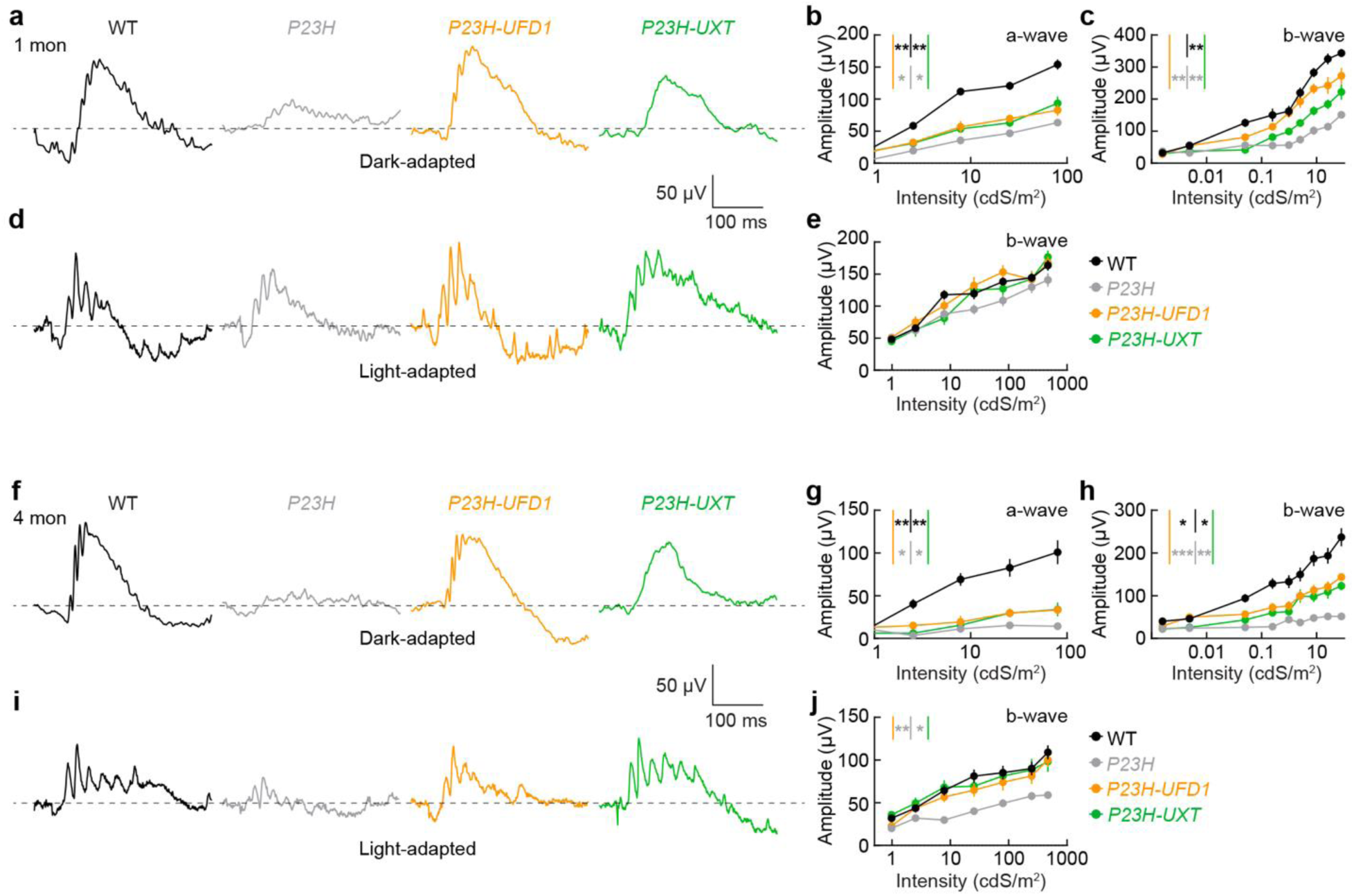
*UFD1* and *UXT* augmentation preserve retinal function in *P23H* mice. **a,f,** Representative dark-adapted (scotopic) ERG traces recorded at one month (**a**) or four months (**f**) in WT (black), *P23H* (gray), *P23H-UFD1* (orange), and *P23H-UXT* (green) mice, shown for flash intensities of 9.83 × 10^−1^ cd·s·m^−2^. **b,c,g,h,** Intensity–response curves for a-wave (**b,g**) and b-wave (**c,h**) amplitudes under dark-adapted conditions at one and four months, respectively. **d,i,** Representative light-adapted (photopic) ERG traces at one (**d**) and four months (**i**) for flash intensities of 470.28 cd·s·m^−2^. **e,j,** Intensity– response curves for the photopic b-wave at 1 (**e**) and 4 months (**j**). Data in (**b,c,e,g,h,j**) are mean ± SEM, with n = 7–15 retinas from 5–9 mice (one-month data) or n = 9–20 retinas from 5–11 mice (four-month data). For all comparisons, black asterisks (*p < 0.05, **p < 0.01) indicate significance vs. WT, while gray asterisks (*p < 0.05, **p < 0.01, ***p < 0.001) indicate significance vs. *P23H* by Monte Carlo permutation testing. Absence of an asterisk denotes p ≥ 0.05.

By four months, rod-driven ERG responses in untreated *P23H* mice had largely disappeared (Fig. 4f–h) ^33,43^. Overexpression of *UFD1* or *UXT* conferred partial rescue of both a- and b-wave amplitudes, with a somewhat stronger effect on the b-wave (Fig. 4f–h), likely reflecting the combined benefit of maintaining rod outer segments and preserving rod–bipolar synapses. At this later time point, cone-driven ERG responses in untreated *P23H* mice were also markedly reduced (Fig. 4i,j). Strikingly, *UFD1* or *UXT* overexpression maintained cone-driven amplitudes at WT levels at four months (Fig. 4i,j).

Together, these findings demonstrate that *UFD1* and *UXT* gene therapies afford robust, long-term functional protection in *P23H* retinas, preventing secondary cone dysfunction and preserving light-evoked responses despite ongoing disease. This highlights the potential for *UFD1* and *UXT* gene augmentation to slow rod loss and safeguard cone function—key goals for mitigating vision loss in RP.

### Neuroprotective therapies preserve visual behaviors in *P23H* mice

Our previous findings demonstrate that *UFD1* or *UXT* gene augmentation in *P23H* mice preserves photoreceptor survival, safeguards both rod- and cone-driven retinal function, and helps maintain synaptic integrity. To determine whether these structural and electrophysiological benefits also protect vision at the behavioral level, we used two established visual assays: the pupillary light response (PLR) and the visual cliff test.

The PLR—a fundamental reflex conserved from mice to humans—adjusts the pupil diameter in response to ambient illumination, to protect the retina from excessive light and increase spatial contrast in the retinal image, enhancing visual acuity ^74, 75^. Under low to intermediate light levels, rods drive the PLR; at bright illumination, the intrinsic light responses of melanopsin-expressing retinal ganglion cells dominate ^74, 76, 77^. In two–three-month-old *P23H* mice, PLR sensitivity decreased by approximately tenfold compared to WT littermates (Fig. 5a–d), reflecting diminished rod function. In striking contrast, *P23H* mice treated with either *UFD1* or *UXT* gene therapy exhibited normal PLR sensitivity, indistinguishable from WT controls (Fig. 5a–d).

**Fig. 5.**
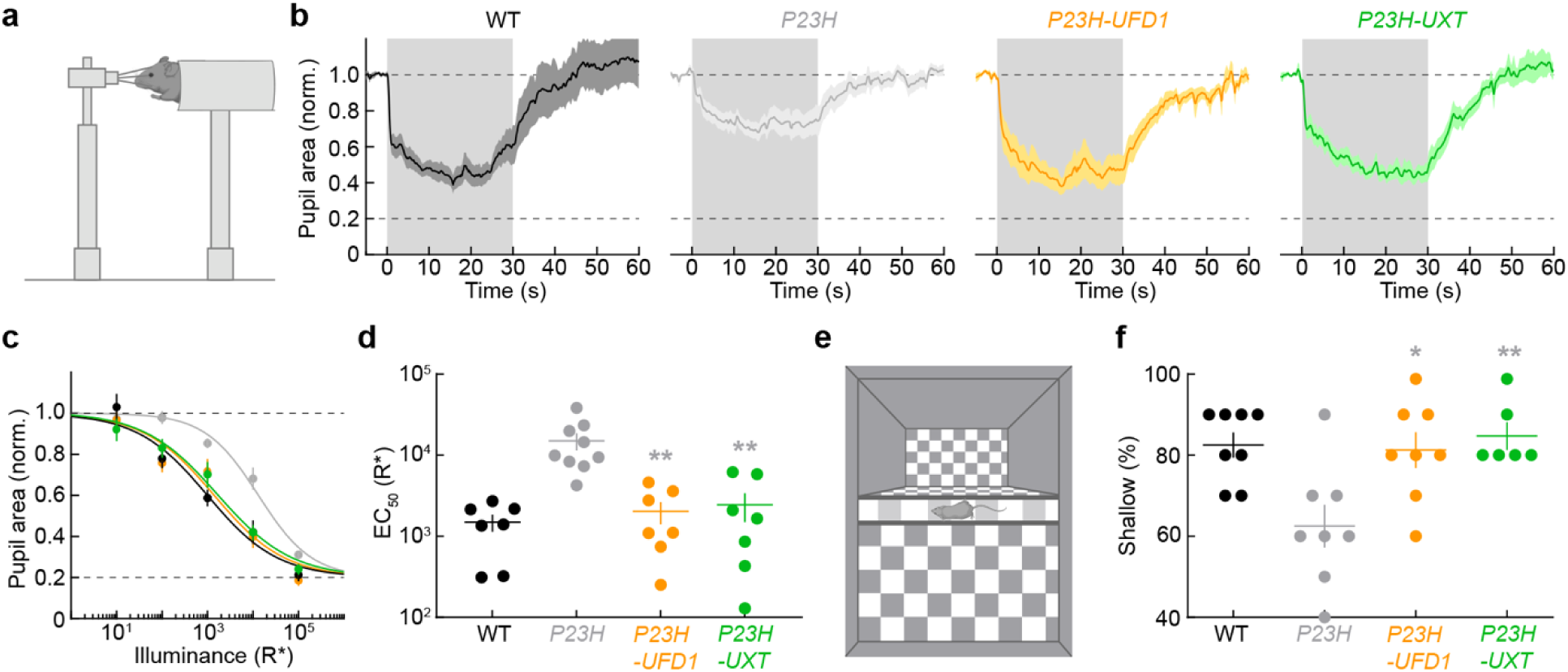
*UFD1* and *UXT* augmentation preserve visual behaviors in *P23H* mice. **a,** Schematic of the apparatus used to measure pupillary light responses (PLRs). **b,** Mean ± SEM pupil area traces from three-month-old WT, *P23H*, *P23H-UFD1*, and *P23H-UXT* mice in response to a light step (gray shading). **c,d,** Intensity–response curves for the PLR (**c**) and half-maximal illuminance (EC_50_) values (**d**). **e,** Schematic of the visual cliff assay, which assesses depth perception based on a preference for stepping onto a shallow (vs. deep) checkerboard pattern. **f,** Percentage of shallow-side choices in two–three-month-old WT, *P23H*, *P23H-UFD1*, and *P23H-UXT* mice. In (**d**) and (**f**), absence of an asterisk indicates p ≥ 0.05, while gray asterisks (*p < 0.05, **p < 0.01) denote significance compared to *P23H* by one-way ANOVA.

We next employed the visual cliff test, commonly used to assess depth perception across species (including human infants) ^78, 79^. Animals on a central ledge choose between stepping down onto a shallow side with the checkered pattern directly beneath it or a deep side where the checkerboard is substantially lower (61 cm in our arena), creating a perceived drop. Conducted under light levels that primarily recruit cones, two– three-month-old *P23H* mice exhibited compromised depth perception and a reduced preference for the shallow side (Fig. 5e,f). Remarkably, *UFD1* or *UXT* overexpression restored their performance to WT levels (Fig. 5e,f).

Together, these results demonstrate that our gene therapies not only protect retinal structure and function but also maintain vision at the behavioral level. By sustaining both rod- and cone-mediated visual behaviors in *P23H* mice, *UFD1* and *UXT* gene augmentation offers a promising therapeutic strategy for mitigating vision loss in RP.

### Human retina explant cultures

A major bottleneck in translation is the transition from animal models to humans. To address this challenge, we cultured explants of adult human retinas to model *P23H* disease and test *UFD1* and *UXT* gene therapies.

We obtained adult human retinas from two sources: (1) organ donors with no history of ocular disease (ischemia times <1 hr) and (2) treatment-naive patients undergoing enucleation for choroidal malignant melanoma (ischemia times <15 min, Supplementary Table 3). Both sources yielded comparable results and were pooled for subsequent analysis. From each retina, we prepared multiple 5 × 5 mm^2^ explants—whose central edge was ∼10 mm from the fovea—and fixed one piece immediately to establish a baseline (Fig. 6a) ^80–82^. Culturing multiple explants in parallel allowed within-donor comparisons and consistent normalization.

**Fig. 6.**
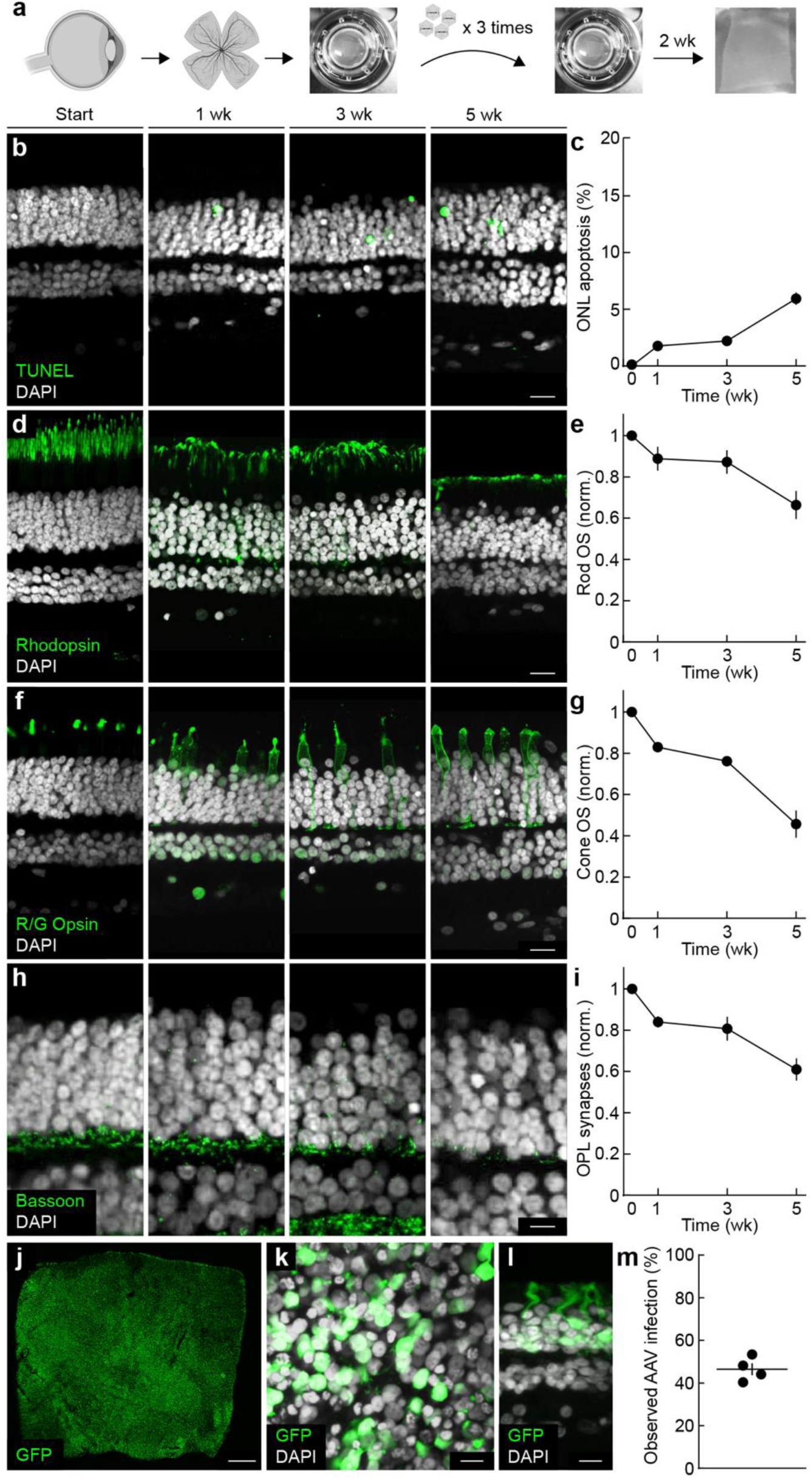
Human retinal explant cultures. **a,** Schematic of the workflow used to establish and maintain adult human retinal explants for up to 5 weeks, with AAV applied at regular intervals. **b,** Representative confocal images of vertical sections at baseline (Start) and after one, three, or five weeks in culture, stained for TUNEL (green) and DAPI (white). Scale bar, 20 µm. **c,** Quantification of TUNEL-positive cells in the ONL from (**b**). **d,** Representative images stained for rhodopsin (green) and DAPI (white), illustrating rod outer segments over time in culture. Scale bar, 20 µm. **e,** Rod outer segment thickness from (**d**), normalized to baseline. **f,** Representative vertical sections stained for red/green opsin (green) and DAPI (white), highlighting cone outer segments. Scale bar, 20 µm. **g,** Cone outer segment thickness from (**f**), normalized to baseline. **h,** Representative sections stained for Bassoon (green) and DAPI (white), labeling presynaptic structures in the OPL. Scale bar, 20 µm. **i,** Bassoon intensity from (**h**), normalized to baseline. In (**c,e,g,i**), data are mean ± SEM; n = 4 donors. **j– l,** Representative examples of explants infected with *AAV-PHP.eB-CAG-GFP* at 1 week, shown in a global whole mount (**j**), a higher-magnification whole mount (**k**), and a vertical section (**l**). Scale bars, 100 µm in (**j**), 20 µm in (**k,l**). **m,** Quantification of AAV infection efficiency in human explants.

TUNEL staining indicated that photoreceptors remained largely viable for up to five weeks, with fewer than 6% of cells in the ONL undergoing apoptosis (Fig. 6b,c). Immunolabeling revealed rod and cone outer segments, along with synapses between photoreceptors and second-order neurons, to be the most sensitive indicators of degeneration, echoing patterns seen in IRDs in vivo ^11, 41^. By three weeks, rod outer segments retained ∼87% of their baseline length (Fig. 6d,e), cone outer segments ∼76% (Fig. 6f,g), and synaptic ribbons ∼80% of baseline staining (Fig. 6h,i). Each measure subsequently declined substantially between three and five weeks (Fig. 6b–i).

We next sought to optimize viral cargo delivery to rod photoreceptors in the explant cultures. We tested different serotypes, promoters, and application protocols. Thus, we found that the *AAV-PHP.eB* serotype ^83^ efficiently infected rods (with limited infection outside the ONL) and that the *CAG* promoter ^84^ drove rapid expression after applying relatively low amounts of virus (3.33 × 10^10^ vg) three times (every other day) at the start of the culture. With this approach, we observed widespread expression of a fluorescent reporter in rods after just one week in culture (Fig. 6j–m).

Together, these findings establish a system in which adult human retinas can be maintained with intact morphology and circuits for at least three weeks, providing a robust platform for modeling retinal degenerations and testing gene therapies, aided by the fast and robust viral manipulation we achieved.

### Disease modeling and neuroprotective gene therapies in the human retina

To recapitulate autosomal dominant RP in adult human retinal explants—derived from donors carrying two wild-type copies of rhodopsin—we delivered a mutant *P23H* rhodopsin transgene via AAV. We verified transgene expression through a myc-tag and incorporated a small-interfering RNA (siRNA) targeting the endogenous wild-type transcript in the 3′UTR to mimic the dominant inheritance pattern (Fig. 7a,b). As a control, we delivered an AAV encoding GFP. In parallel, we introduced our therapeutic constructs for *UFD1* or *UXT*, each bearing a flag-tag for verification (Fig. 7a,b).

**Fig. 7.**
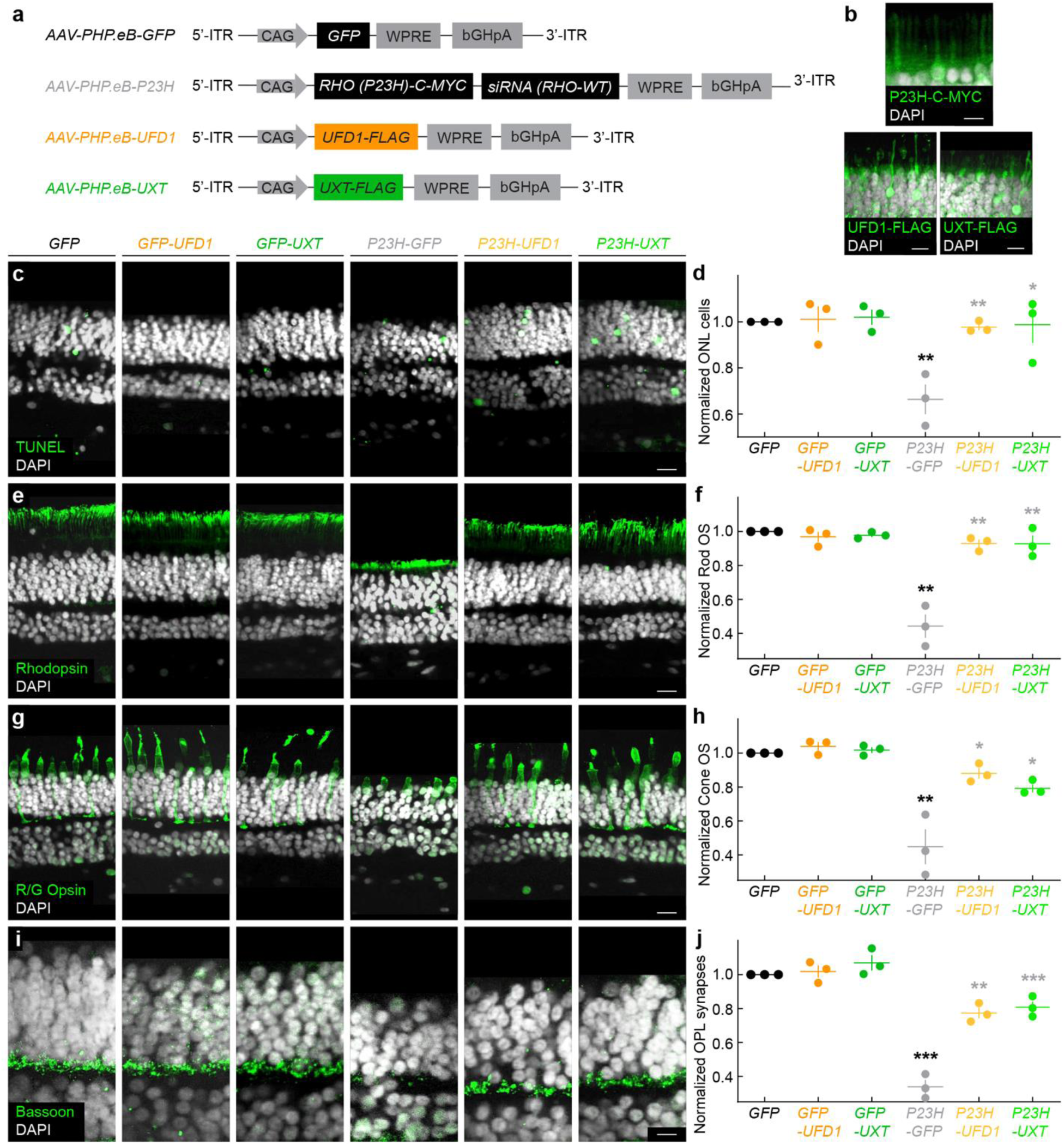
*UFD1* and *UXT* augmentation protect photoreceptors in human *P23H* retinas. **a,** Schematic of the AAV constructs used to introduce the *P23H* rhodopsin transgene (*RHO [P23H]-C-MYC* plus a siRNA targeting wild-type rhodopsin) and to overexpress *UFD1-FLAG* or *UXT-FLAG* in adult human retinal explants. A separate AAV encoding GFP serves as a control. **b,** Representative confocal images confirming *P23H–C-MYC*, *UFD1–FLAG*, and *UXT–FLAG* expression at one week post-infection. Scale bar, 5 μm. **c,** TUNEL (green) and DAPI (white) labeling in vertical sections at three weeks for each experimental group: *AAV-GFP* alone (*GFP*), *GFP + UFD1* (*GFP-UFD1*), *GFP + UXT* (*GFP-UXT*), *P23H + GFP* (*P23H-GFP*), *P23H + UFD1* (*P23H-UFD1*), and *P23H + UXT* (*P23H-UXT*). Scale bar, 20 μm. **d,** Quantification of ONL cell counts in (**c**), normalized to GFP. **e,** Representative sections stained for rhodopsin (green) and DAPI (white). Scale bar, 20 μm. **f,** Rod outer segment (OS) thickness from (**e**), normalized to GFP. **g,** Representative images stained for red/green (R/G) opsin (green) and DAPI (white). Scale bar, 20 μm. **h,** Cone OS thickness from (**g**), normalized to GFP. **i,** Sections stained for Bassoon (green) and DAPI (white), highlighting OPL synapses. Scale bar, 10 μm. **j,** Bassoon intensity from (**i**), normalized to GFP. For (**d,f,h,j**), absence of an asterisk indicates p ≥ 0.05. Black asterisks (**p < 0.01, ***p < 0.001) denote significance vs. GFP; gray asterisks (*p < 0.05, **p < 0.001, ***p < 0.001) denote significance vs. *P23H-GFP* by one-way ANOVA.

Under control (GFP-only) conditions, overexpression of *UFD1* or *UXT* did not alter photoreceptor survival (Fig. 7c,d). However, expression of *P23H* rhodopsin drove significant rod loss by three weeks in culture, which was rescued to near-control levels by *UFD1* or *UXT* overexpression (Fig. 7c,d). Consistent with these findings, rod outer segments—otherwise severely compromised by *P23H* rhodopsin—were also preserved by both *UFD1* and *UXT* therapy (Fig. 7e,f).

Although rod photoreceptors degenerate first in RP, it is the subsequent loss of cone function—critical for bright-light and high-acuity vision—that most profoundly compromises patients’ everyday visual tasks in modern, well-lit environments. In our explant model, *P23H* rhodopsin triggered secondary cone outer segment loss by three weeks, and this degeneration was effectively prevented by *UFD1* or *UXT* overexpression (Fig. 7g,h), recapitulating our observations in mice (Fig. 3–5). Furthermore, *P23H* rhodopsin induced synaptic pathology in the outer plexiform layer, mirroring early synapse loss observed in IRDs and other neurodegenerative diseases (Fig. 7i,j) ^11, 41, 72, 73^. Again, co-expression of *UFD1* or *UXT* prevented these synaptic deficits (Fig. 7i,j).

In summary, these results establish an experimentally accessible model of *P23H* disease in adult human retina explants and demonstrate that *UFD1* and *UXT* gene therapies—originally developed in mice—can also protect human photoreceptors. These findings highlight the translational potential of *UFD1* and *UXT* augmentation for preserving vision in patients with autosomal dominant RP and related IRDs.

## Discussion

In this study, we (1) performed a genome-wide in vivo CRISPR screen in a *P23H* mouse model of RP to systematically identify neuroprotective genes in rod photoreceptors; (2) discovered a set of gene knockouts that accelerated rod degeneration in *P23H* retinas, (3) validated top-ranked candidates in a secondary screen, and (4) confirmed that none were essential in the absence of the *P23H* allele, highlighting their disease-specific roles; (5) showed that overexpressing two of these genes—*UFD1* and *UXT*—preserves photoreceptor structure, synaptic integrity, and rod- and cone-driven visual function in *P23H* mice; (6) developed a human *P23H* model by expressing mutant rhodopsin in adult human retinal explants, enabling direct assessment of neuroprotective therapies in a clinically relevant context; and (7) demonstrated that *UFD1* and *UXT* augmentation likewise safeguards rods, cones, and synapses in the human retina, underscoring their translational potential for autosomal dominant RP and related IRDs.

Most previous efforts to identify neuroprotective strategies—both in the retina and in the broader nervous system—have focused on testing individual, biologically plausible mechanisms ^18, 19^. While this hypothesis-driven approach has led to some successes, it has inherently limited the scope of exploration and left large portions of the potential therapeutic landscape unexplored. By contrast, our genome-wide CRISPR screen systematically evaluated the entire protein-coding transcriptome, enabling an unbiased and comprehensive search for neuroprotective factors. This approach not only identified well-characterized genes with known roles in retinal health (e.g., *Hgsnat*, *Prpf4*, *Bbs1*) ^85–87^ but also uncovered relatively unstudied genes among the top hits (Fig. 1; Supplementary Tables 1 and 2). By directly ranking genes based on their impact on rod photoreceptor viability, our screen prioritizes molecular pathways with high therapeutic potential, creating a roadmap for future mechanistic studies and drug development efforts.

Genome-wide CRISPR screens have identified novel therapeutic targets in cancer ^20–22^, autoimmune disorders ^23, 24^, and infectious diseases ^25–27^. However, their in vivo application to the nervous system has been limited for several reasons ^29^: neurons are nondividing cells with lower CRISPR editing efficiency ^88, 89^; it can be challenging to infect enough neurons with pooled libraries at the appropriate multiplicity ^90, 91^; and the cell-type diversity and type-specific susceptibility of neurons in many brain areas complicate the interpretation of screening data ^92–94^. We overcame some of these challenges by focusing on rod photoreceptors—a single, abundant neuronal type in the retina ^93, 95^—that can be efficiently transduced in vivo through subretinal lentiviral injections (Fig. 1). We demonstrate the feasibility of a genome-wide in vivo CRISPR screen in rods of *P23H* mice, which model the most common cause of autosomal dominant RP in the United States ^4, 30^. Future CRISPR screens in other IRD models could help determine which neuroprotective mechanisms are pathway-specific and which act broadly to enhance rod survival ^11^.

Rhodopsin is folded in the ER on its way to the outer segment disc membrane, but the *P23H* mutation disrupts this process. The resulting protein misfolding triggers ER stress and the formation of rhodopsin aggregates, causing photoreceptor degeneration ^96–100^. Cells maintain proteostasis through multiple quality-control pathways, including autophagy and proteasomal degradation ^101^. Our CRISPR screen highlighted vacuolar ATPases (V-ATPases) as the most strongly depleted gene family (Extended Data Fig. 2c,d); these proton pumps drive lysosomal acidification and are essential for degrading misfolded proteins ^102, 103^. When lysosomal acidification is disrupted—as in the V-ATPase knockouts in our screen—*P23H* rhodopsin may accumulate faster, exacerbating stress and accelerating cell death. In line with this idea, inhibiting V-ATPases with bafilomycin A1 doubled *P23H* rhodopsin levels in vitro ^60^, supporting the critical role of lysosomal acidification in autophagic degradation. Future experiments should explore whether enhancing V-ATPase activity (for example, via gene augmentation) can mitigate photoreceptor degeneration in *P23H* retinas and related IRDs.

Aggrephagy refers to the selective autophagy of protein aggregates ^104, 105^. Misfolded rhodopsin, including the *P23H* variant, is prone to aggregate in photoreceptors ^96, 99^. UXT links aggregates to p62, a key autophagy receptor that sequesters aggregates in condensates and targets them to phagophores for lysosomal breakdown ^68^. Consistent with the neuroprotective role we identified, *UXT* knockout mice exhibit features of RP ^65^. Our findings suggest that *UXT* overexpression promotes aggregate clearance and preserves photoreceptor survival, morphology, and connectivity in both mouse and human *P23H* models, although the mechanisms by which *UXT* stabilizes or removes toxic rhodopsin species require further study.

Another crucial pathway for degrading aberrant rhodopsin involves the proteasome. Misfolded ER proteins such as *P23H* rhodopsin are tagged with ubiquitin and channeled to the proteasome through ER-associated degradation (ERAD) ^60, 100, 106^. In this process, the p97/VCP AAA-ATPase extracts ubiquitinated substrates from the ER membrane; UFD1, an essential p97/VCP cofactor, recognizes these tagged proteins and helps deliver them for proteasomal destruction ^62, 107^. In vitro work has shown that *P23H* rhodopsin undergoes ubiquitination ^108^, with p97/VCP promoting its dislocation and degradation ^100, 109^. We hypothesize that *UFD1* overexpression augments this ERAD pathway and mitigates the toxic buildup of misfolded *P23H* rhodopsin in photoreceptors, contributing to improved visual outcomes in mice and, potentially, humans. Direct experimental tests of this mechanism remain a key next step.

Together, these findings indicate that *UXT* and *UFD1* may enhance the clearance of *P23H* rhodopsin through distinct yet complementary mechanisms, suggesting that dual gene augmentation might offer greater therapeutic benefit than either approach alone ^110^. It will be important to examine this possibility, alongside strategies to elevate proteasome function ^111^ and limit ER stress ^112^, to determine whether boosting multiple proteostasis pathways confers synergistic neuroprotection. Moreover, because protein misfolding and aggregation are central to numerous degenerative disorders—including Alzheimer’s ^12–14^, Huntington’s ^15, 16^, and amyotrophic lateral sclerosis ^14, 17^—it will be intriguing to test whether the *UXT* and *UFD1* strategies developed here can be adapted for neurodegenerative conditions beyond the retina.

A major challenge in translational research is bridging the gap between animal models and human tissues, a transition where many promising strategies fail. Although mouse and human rod photoreceptors share fundamental features, they also differ in their transcriptomes ^80, 92, 93, 113, 114^, morphology ^115^, metabolism ^116^, and connectivity ^117^. These differences limit the predictive power of mouse models for RP therapies, underscoring the importance of testing candidate interventions in human retinal systems.

Retinal organoids derived from human pluripotent stem cells have been transformative for modeling early stages of retinal development and congenital disorders ^37, 38, 80, 118^. However, organoids face limitations in recapitulating adult-onset diseases: the reprogramming step resets the age signatures of donor cells to a fetal state ^119–121^, and even after many months in culture, most organoid-derived photoreceptors remain functionally immature ^122^, with rudimentary synapses ^80, 118^. Furthermore, the early degeneration of ganglion cells disrupts downstream circuits, compromising the organoid’s ability to reflect mature retinal architecture and disease progression ^118^.

Adult human retinal explants preserve the mature organization and connectivity of photoreceptors and downstream circuits, making them uniquely suited to study late-onset retinal degenerations and to test gene therapy approaches ^35, 123^. Although they have proven valuable for refining viral transduction strategies ^81^, improving gene editing efficiencies ^124^, and exploring potential treatments to restore light responses ^82^, explants from donors do not typically carry pathogenic mutations. To address this limitation, we used an AAV-based approach to simultaneously express a human *P23H* rhodopsin transgene and knock down the endogenous WT allele, thereby creating an experimentally accessible adult human retinal model of *P23H* disease. This model reproduced key features of in vivo photoreceptor degeneration and enabled us to validate the translational potential of *UFD1* and *UXT* gene augmentation therapies developed in mice (Fig. 7). Future efforts could employ similar methods to model other gain-of-function IRDs or utilize CRISPR-mediated knockouts to recreate loss-of-function mutations, ultimately broadening the scope of human retinal disease modeling and therapy testing.

## Methods

### Human tissue samples

Human retina tissue was obtained from multi-organ donors by sampling non-transplantable eye tissue that was removed during cornea harvesting for transplantation. Donors with a documented history of eye disease were excluded from this study. Personal identifiers were removed, and samples were anonymized before processing. Additional human retina tissue was obtained from treatment-naive patients enucleated for choroidal malignant melanomas (Supplementary Table 3). The research protocol for this study was approved by the Washington University School of Medicine Human Research Protection Office and the Institutional Review Board (IRB, Protocol #201805087) in compliance with HIPAA guidelines and the tenets of the Declaration of Helsinki. Written informed consent was obtained from the choroidal malignant melanoma patients prior to inclusion in the study.

### Mice

For the CRISPR screen, we generated *P23H Nrl-Cas9* mice. We first bred a homozygous *P23H* knockin line (RRID:IMSR_JAX:017628) ^33^ with an *Nrl-Cre* transgenic line, in which Cre recombinase is driven by a rod progenitor–specific promoter (RRID:IMSR_JAX:028941) ^125^. The resulting Cre-positive offspring were then bred to a knockin line expressing Cas9 from a ubiquitously active locus in a Cre-dependent manner (RRID:IMSR_JAX:026175) ^126^. For negative and positive selection in our screen, *P23H* heterozygous Cre-positive mice were used, while their Cre-negative littermates served as library input controls. For candidate overexpression experiments, the homozygous *P23H* knockin line was bred to the *Nrl-Cre* line to generate *P23H Nrl-Cre* mice. Cre-positive animals were used to overexpress either *UFD1* or *UXT*, whereas Cre-negative littermates injected with AAVs served as controls. All mice in this study were backcrossed for more than five generations onto a C57Bl6/J background (RRID:IMSR_JAX:000664). For pupillary light reflex (PLR) testing, head plates were affixed to the skull using dental resin (Parkell) in mice anesthetized with ketamine (0.1 mg/g body weight) and xylazine (0.01 mg/g body weight). After surgery, mice were allowed to recover for at least one week prior to PLR testing. All procedures were approved by the Animal Studies Committee of Washington University School of Medicine (Protocol #23-0116) and performed in accordance with the National Institutes of Health *Guide for the Care and Use of Laboratory Animals*. We observed no sex-specific differences in our results; therefore, data from both male and female mice were pooled in analyses.

### Genome-wide in vivo CRISPR screen

#### CRISPR library preparation

The Mouse Brie CRISPR knockout pooled library was a gift from David Root and John Doench (Addgene #73633) ^44^. It contains 78,637 sgRNAs targeting 19,674 genes, covering the protein-coding mouse genome. Each sgRNA is expressed under the control of a U6 promoter. The plasmid DNA for the library was amplified to achieve >1,000× coverage following the recommended protocol. To confirm proper representation and distribution of sgRNAs, next-generation sequencing was performed on the purified plasmid DNA prior to lentivirus production. The VSV-G–pseudotyped lentiviral library was packaged by SignaGen and achieved a final titer of 1.26 × 10^9^ TU/mL for in vivo delivery.

#### Subretinal LV injection

For subretinal LV injections, *P23H Nrl-Cas9* mouse pups (P0–1) were anesthetized on ice. A total of 210 nL of either the LV library (1.26 × 10^9^ TU/mL) or *LV-EF1α-GFP* (1 × 10^9^ TU/mL, Hope Center Viral Vectors Core at Washington University School of Medicine) was injected into the subretinal space using a Nanoject II injector (Drummond).

#### Estimating LV multiplicity of infection

We assume that LV infection of photoreceptors follows a Poisson distribution and denote the average number of lentiviruses infecting a cell as 𝑚 (i.e., the multiplicity of infection). According to the Poisson distribution, the probability that a given cell gets infected by exactly 𝑘 viruses is:

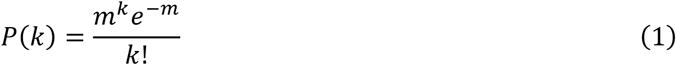

The observed labeling fraction 𝑓 (i.e., the fraction of photoreceptors expressing GFP, Fig. 1d–g) is the probability that a cell is infected by at least one virus. This probability is:

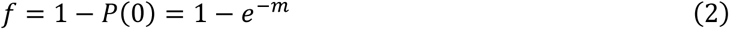

Solving for 𝑚:

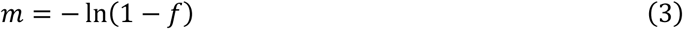

Substituting this in the Poisson formula:

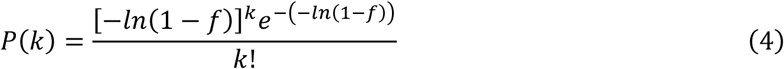

Notice that:

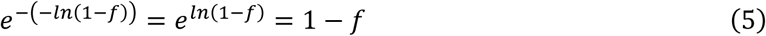

Thus, for a given observed labeling fraction 𝑓, the probability that a photoreceptor is infected by exactly 𝑘 LVs is:

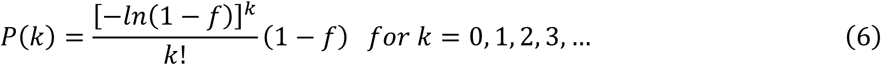

#### Retinal harvest and genomic DNA isolation

At P30, mice were euthanized by CO₂, and eyes were enucleated. Individual retinas were immediately isolated and lysed overnight at 56°C in a homemade lysis buffer containing 50 mM Tris, 1 mM CaCl₂, 0.1% Tween-20, and 10 U/mL proteinase K (New England Biolabs, CAT#P8107S). Lysates were centrifuged at 14,000 rpm at room temperature to remove debris, and DNA was precipitated with isopropanol, washed with 70% ethanol, and dissolved in ddH₂O. Each retina served as an independent sample for downstream analysis. Genomic DNA samples with an A₂₆₀/A₂₈₀ ratio below 1.6 were excluded.

#### LV integration testing

To identify and exclude samples exhibiting insufficient LV integration, genomic DNA (500 ng) from each sample was subjected to PCR amplification targeting the lentiviral packaging region. The following primer pair was utilized:

LV-Forward: ACC TGA AAG CGA AAG GGA AAC

LV-Reverse: CAC CCA TCT CTC TCC TTC TAG CC

PCR products were subsequently analyzed by agarose gel electrophoresis. DNA extracted from retinas infected with *LV-EF1α-GFP*, with previously established transduction efficiency (as determined by cell counting, Fig. 1d–g), served as a reference control. Only samples yielding PCR amplification comparable to the positive control were included in subsequent experiments.

#### sgRNA PCR amplification and sequencing

sgRNAs were amplified from genomic DNA by nested PCR. For the first PCR (PCR_1_), all DNA from each sample was divided among multiple 50 μL reactions (each containing up to 50 μg DNA) using Q5 High-Fidelity DNA Polymerase (New England Biolabs, CAT#M0491). The PCR_1_ reaction contained an equimolar mix of the following phased forward primers:

ACGACGCTCTTCCGATCTTTGTGGAAAGGACGAAACACCG ACGACGCTCTTCCGATCTCTTGTGGAAAGGACGAAACACCG ACGACGCTCTTCCGATCTGCTTGTGGAAAGGACGAAACACCG ACGACGCTCTTCCGATCTAGCTTGTGGAAAGGACGAAACACCG

ACGACGCTCTTCCGATCTCAACTTGTGGAAAGGACGAAACACCG ACGACGCTCTTCCGATCTTGCACCTTGTGGAAAGGACGAAACACCG ACGACGCTCTTCCGATCTACGCAACTTGTGGAAAGGACGAAACACCG ACGACGCTCTTCCGATCTGAAGACCCTTGTGGAAAGGACGAAACACCG and a single reverse primer:

### GTGACTGGAGTTCAGACGTGTGCTCTTCCGATCTTCTACTATTCTTTCCCCTGCACTGT

PCR_1_ cycling conditions were as follows: 98°C for 30 s; 8 cycles of 98°C for 10 s, 55°C for 30 s, 72°C for 30 s; 10 cycles of 98°C for 10 s and 72°C for 60 s; followed by a final extension at 72°C for 5 min and a hold at 4°C. The PCR_1_ products were purified using SeraPure DNA isolation beads (Sera-mag SpeedBeads [Cytiva], prepared according to ^127^) at a 1:1 reaction:bead ratio, washed twice with 80% ethanol, and eluted in 25 μL ddH₂O.

The eluted DNA from each sample was pooled and thoroughly mixed. A 2.5 μL aliquot of this pooled DNA was used as the template for a second PCR (PCR_2_) using Q5 High-Fidelity DNA Polymerase and barcoded primers that incorporated Illumina sequencing adapters. The primer sequences used for PCR_2_ were:

PCR_2_-forward: AATGATACGGCGACCACCGAGATCTACAC-[10bp-index]-ACACTCTTTCCCTACACGACGCTCTTCCGATCT

PCR_2_-reverse: CAAGCAGAAGACGGCATACGAGAT-[10bp-index]-GTGACTGGAGTTCAGACGTG

PCR_2_ cycling conditions were 98°C for 30 s; 8 cycles of 98°C for 10 s, 66°C for 30 s, 72°C for 30 s; 5 cycles of 98°C for 10 s and 72°C for 60 s; followed by a final extension at 72°C for 5 min and a hold at 4°C. The PCR_2_ products were then purified using SeraPure beads (1:1 reaction:bead ratio), washed twice with 80% ethanol, and eluted in 30 μL ddH_2_O.

All final PCR products were verified for size using a Bioanalyzer High Sensitivity DNA chip and quantified by Qubit before submission for sequencing at the Washington University Genome Technology Access Center and the McDonnell Genome Institute. Libraries were sequenced on the Illumina NovaSeq X platform with 150 bp paired-end reads.

#### Screen Analysis

Raw sequencing data were trimmed using Cutadapt (v0.6.7) ^128^. Reads matching each gRNA were quantified with custom scripts and then analyzed using drugZ ^45^. Enrichment of specific categories was assessed using DAVID ^129^ and Gene Set Enrichment Analysis (GSEA, v4.3.3) ^130^.

### RNA-seq reanalysis and integration

Raw RNA-seq data for sorted mouse rod photoreceptors (SRR3662498, SRR3662513, SRR3662514) ^56^ and human peripheral retina (SRR5591599) ^57^ were obtained from the SRA. Reads were trimmed with Trim Galore! (v0.6.7), a wrapper for Cutadapt (v3.4) and FastQC (v0.11.9), then mapped to the GRCm38 (mouse) or GRCh38 (human) genome builds using STAR (v2.7.0) ^131^. The mapped reads were processed with Samtools (v1.9.4) ^132^, and gene counts were generated using HTseq (v0.11.2) ^133^. Additionally, count matrices from a single-cell RNA-seq dataset profiling the whole mouse retina (GSE63473) ^93^ were downloaded. All data were analyzed using Monocle 3 (v0.1.3) ^134–137^, and relative normalized expression values were calculated for each cell type. Final plots were generated with gplots (v3.2.0) and ggplot2 (v3.5.1).

### Validation of the CRISPR screen

To validate the individual effects of top-depleted hits on photoreceptor survival, sgRNAs targeting each candidate gene were designed using the VBC Score algorithm. Two of the highest-ranked sgRNAs— distinct from those used in the original CRISPR screen library—were cloned into a vector Cre-dependently expressing tdTomato from a CAG promoter and each sgRNA from a U6 promoter (*pCAG-Flex-tdTomato-U6-sgRNA*). Constructs were delivered to rod photoreceptors via in vivo electroporation. Briefly, mouse pups at postnatal day 0–1 (P0–P1) were anesthetized on ice, and 210 nL of *pCAG-Flex-tdTomato-U6-sgRNA* (3µg/µl) were injected into the subretinal space using a Nanoject II injector (Drummond). Five 80 V pulses, each 50 ms in duration, were generated by an ECM 830 electroporator (BTX, Harvard Apparatus) and delivered using tweezer electrodes, with the anode placed on the injected eye ^58, 59, 138^. Mice electroporated with a construct including a non-targeting sgRNA served as controls.

### Gene augmentation therapies

#### Construct generation and AAV production

For candidate overexpression, open reading frames (ORFs) of human UFD1 and UXT were obtained from GenScript, inverted, and cloned into the *pAAV-CAG-Flex-Twitch2B* vector, replacing the Twitch2B sequence. AAVs were then packaged with the *AAV-PHP.eB* capsid by SignaGen Laboratories, achieving titers exceeding 1 × 10^13^ vector genomes (vg) per milliliter.

#### Subretinal AAV injection

For subretinal AAV injections, *P23H Nrl-Cre* mouse pups (P2) were briefly anesthetized on ice. A total of 280 nL of either *AAV-Flex-UFD1* (*UFD1* ORF NM_005659.6) or *AAV-Flex-UXT* (UXT ORF NM_153477.2) was delivered into the subretinal space using a Nanoject II injector (Drummond).

### Electroretinograms (ERGs)

ERGs were recorded from one- and four-month-old mice as previously described. Briefly, mice were dark-adapted overnight, anesthetized with ketamine (0.1 mg/g body weight) and xylazine (0.01 mg/g body weight), and their pupils were dilated with 1% atropine sulfate (Falcon Pharmaceuticals). Recording electrodes embedded in contact lenses were placed over both corneas, and the mice were maintained at 37 ± 0.5°C throughout the recordings using a heating pad and rectal temperature probe (FHC, Bowdoin, ME, USA). White light flashes (<5 ms) were delivered, and responses from *WT*, *P23H*, *P23H-UFD1*, and *P23H-UXT* mice (*P23H-UFD1* and *-UXT* refer to *P23H Nrl-Cre* mice injected with *AAV-Flex-UFD1* and *-UXT*, respectively) were acquired using the UTAS Visual Electrodiagnostic Testing System (LKC Technologies). Four to ten responses were averaged at each light level. The a-wave amplitude was measured as the difference between the minimum response within the first 50 ms after flash onset and the baseline voltage at flash onset. The b-wave amplitude was measured as the difference between the low-pass filtered (15–25 Hz) b-wave peak and the a-wave amplitude. All ERG analyses were performed using custom scripts written in MATLAB.

### Visual behaviors

#### Pupillary Light Reflex (PLR)

PLR testing was performed as previously described ^74, 139^. Briefly, the left eye of head-fixed, awake mice was tracked under infrared illumination using an ETL-200 eye-tracking system (ISCAN). Stimuli were simultaneously presented to the right eye via an Arduino-controlled 525 nm LED (Thorlabs), with intensity adjusted through a series of neutral density (ND) filters (Thorlabs). Illuminance steps ranged from 10^0^ to 10^5^ R* in half-log (10^0.5^) increments. Each step was preceded by 30 s of darkness and followed by 30 s of post-illumination recovery to baseline; stimuli were separated by at least 2 min in darkness. Pupil constriction was normalized to the fully dark-adapted pupil area. The relative pupil area at each light level was calculated by averaging the pupil size over a 5 s interval at the point of maximal constriction. EC_50_ values, representing the light intensity required to achieve 50% of maximal pupil constriction, were determined by fitting data to a Hill equation model.

#### Visual Cliff

Visual cliff testing was conducted on a 56 × 41 cm platform with a 3.8 × 1.7 cm (height × width) ridge positioned across its center. On one side of the ridge, a checkerboard pattern was placed immediately beneath the platform (the shallow side), whereas on the other side, the same checkerboard pattern was located 61 cm below the platform (the deep side) ^140^. Each mouse underwent ten trials, with the orientation of the shallow and deep sides randomly alternated between trials. The preference for stepping onto the shallow or deep side was recorded in each trial.

### Human retinal explant culture

Human retinal explant cultures were established as previously reported ^81, 82^. Posterior poles (globes without the cornea) were collected from organ donors with an ischemia time of less than one hour, and the retinas were isolated from the sclera and pigment epithelium ^80^. Alternatively, globes from patients undergoing enucleation surgery for choroidal malignant melanomas were bisected under transillumination within 10 min of central artery occlusion on the non-tumor side of the optic nerve. Retinas were transported from the tissue donor site and operating rooms to the laboratory (10-30 min) in dark container filled with continuously oxygenated HEPES-buffered Ames’ medium. Retinas were then dissected into 5 × 5 mm pieces and placed on 6-well polycarbonate membrane transwells (Corning) with the ONL facing upward. Explants with the same eccentricity were used for comparison. Retinal explants were cultured at 37°C in 5% CO₂ for three weeks in DMEM/F-12 nutrient medium (ThermoFisher), supplemented with 0.1% BSA, 10 μM O-acetyl-l-carnitine hydrochloride, 1 mM fumaric acid, 0.5 mM galactose, 1 mM glucose, 0.5 mM glycine, 10 mM HEPES, 0.05 mM mannose, 13 mM sodium bicarbonate, 3 mM taurine, 0.1 mM putrescine dihydrochloride, 0.35 μM retinol, 0.3 μM retinyl acetate, 0.2 μM (±)-α-tocopherol, 0.5 mM ascorbic acid, 0.05 μM sodium selenite, 0.02 μM hydrocortisone, 0.02 μM progesterone, 1 μM insulin, 0.003 μM 3,3′,5′-triiodo-l-thyronine, 2,000 U penicillin, and 2 mg streptomycin (Sigma-Aldrich), following the protocol described by ^81^. The culture medium was changed every other day.

### Disease modeling and gene therapies in human retinal explants

#### Construct generation and AAV production

For *P23H* rhodopsin overexpression and WT rhodopsin knockdown, the open reading frame (ORF) of human rhodopsin (purchased from GeneCopoeia) was tagged with C-MYC and cloned into the pLKO.1 puro vector. The *P23H* mutation was introduced by PCR using the following forward primer:

GTGGTACGCAGCCACTTCGAGTACCCACAGTAC

and reverse primer:

TGGGTACTCGAAGTGGCTGCGTACCACACCCGT

A small interfering RNA (siRNA) targeting the 3′ UTR of WT rhodopsin (sequence: AGGACTCTGTGGCCGACTATA) was designed using the GPP Web Portal and embedded in the 3′ region of *rhodopsin-P23H-C-MYC* as previously described ^141^.

For candidate overexpression, the ORFs of human *UFD1* and *UXT* (purchased from GenScript) were tagged with FLAG and cloned into the *AAV-CAG-GFP* construct to replace GFP. All AAVs were packaged with an *AAV-PHP.eB* capsid by SignaGen Laboratories at titers exceeding 1 × 10^13^ vg mL.

#### AAV administration

For AAV infection, 3.33 × 10^10^ vg of each AAV was added to the cultured tissue every other day for a total of three applications (1 × 10^11^ vg in total) at the start of the culture period. Tissues were fixed at three days and again at two weeks after the final viral treatment, for expression confirmation and assessment of downstream effects, respectively. To model *P23H* disease in human retinal explants, the same amount of control (*GFP*), model (*P23H*), or rescue (*UFD1* and *UXT*) AAVs was used for the relevant treatment groups.

### Tissue preparation and immunohistochemistry

#### Mouse retinal dissection

Mice were euthanized with CO₂ and enucleated. Retinas were isolated in HEPES-buffered mouse artificial cerebrospinal fluid (mACSF-HEPES) composed of 119 mM NaCl, 2.5 mM KCl, 2.5 mM CaCl₂, 1.3 mM MgCl₂, 1 mM NaH₂PO₄, 11 mM glucose, and 20 mM HEPES (pH 7.37, adjusted with NaOH). The isolated retinas were flat-mounted on membrane disks (HABGO1300, Millipore) and fixed for 30 min in 4% paraformaldehyde (PFA).

#### Human retinal explant preparation

Retinal explants were removed from the transwell membrane in phosphate-buffered saline (PBS) and fixed for 30 min in 4% PFA.

#### Flat-mount staining

Retinas were cryoprotected by sequential incubation in 10% sucrose in PBS for 1 hr at room temperature (RT), 20% sucrose in PBS for 1 hr at RT, and 30% sucrose in PBS overnight at 4°C. After three freeze-thaw cycles, retinas were washed three times for 10 min each in PBS at RT, then blocked in 10% normal donkey serum (NDS) in PBS for 2 hr at RT. Next, they were incubated with primary antibodies (chicken anti-GFP, ThermoFisher, RRID:AB_2534023; mouse anti-Rhodopsin, MilliporeSigma, RRID:AB_260838; rabbit anti-cone arrestin, MilliporeSigma, RRID:AB_1163387; rabbit anti-R/G Opsin, MilliporeSigma, RRID:AB_177456, mouse anti-CtBP2, BD Biosciences, RRID:AB_399431; mouse anti-Bassoon, ThermoFisher, RRID:AB_2066981; sheep anti-mGluR6 ^142^; mouse anti-PKCɑ, ThermoFisher, RRID:AB_477375) in 5% NDS and 0.5% Triton X-100 in PBS for 5 days at 4°C. Retinas were then washed three times for 1 hr each in PBS at RT, incubated overnight at 4°C with the appropriate secondary antibodies (Donkey anti-chicken IgY Alexa 488, ThermoFisher, RRID:AB_2534096; Donkey anti-rabbit IgG Alexa 488, ThermoFisher, RRID:AB_2556546; Donkey anti-mouse IgG Alexa 568, ThermoFisher, RRID:AB_2534013; Donkey anti-sheep IgG Alexa 633, ThermoFisher, RRID:AB_2535754) along with DAPI to stain nuclei, washed again three times for 1 hr at RT in PBS, and mounted in Vectashield medium (Vector Laboratories, RRID:AB_2336789).

#### Slice staining

Retinal cups were embedded in 4% agarose and cut into 50 μm sections using a vibratome (VT1000A, Leica). Sections were blocked for 2 hr at RT in 10% NDS in PBS, then incubated overnight at 4°C with primary antibodies (see Flat-mount staining) in 5% NDS and 0.5% Triton X-100 in PBS. Afterward, slices were washed in PBS three times for 10 min each at RT, stained with secondary antibodies (see Flat-mount staining) and DAPI for 2 hr at RT, washed again three times for 10 min in PBS at RT, and finally mounted in Vectashield medium.

### Imaging and analysis

Confocal images were acquired using an FV1000 laser scanning microscope (Olympus) equipped with a 60×/1.35 NA oil-immersion objective. To assess viral infection, tiling images were obtained on an LSM 800 microscope (Zeiss). Images were processed and analyzed with Fiji ^143^. Nuclei count, apoptotic cell count, ONL thickness, rod outer segment thickness, cone outer segement thickness, and outer plexiform layer synapse intensity were measured in 211.97 × 211.97 µm² images, while rod bipolar cell (RBC) mGluR6 puncta were quantified in 70.66 × 70.66 µm² images. Consistent imaging parameters were applied for all outer plexiform layer synapse intensity measurements. Three to five images per animal were averaged for each comparison.

### RT-PCR

Mouse retinas were dissected and flash-frozen in liquid nitrogen. Total RNA was isolated using the RNeasy Micro Kit (Qiagen, CAT#74004) and reverse-transcribed into cDNA with SuperScript IV Reverse Transcriptase (ThermoFisher, CAT#18090010). Cre-dependent inversions of *UFD1* and *UXT* were detected by PCR with the following primers:

*UFD1-Forward:* GTGGAGAGCGTCAACCTTCA

*UXT-Forward:* TGGCCACCTACTCCAAATTC

*Reverse:* CCGAAGGGACGTAGCAGA

A housekeeping gene, Lactate dehydrogenase A (LDHA), served as an internal reference using:

*LDHA-Forward:* CTGAGCACACCCATGTGAGA

*LDHA-Reverse:* AGCAACACTCCAAGTCAGGA

PCR products were analyzed by agarose gel electrophoresis.

### Statistics

Statistical differences were evaluated using one-way or two-way ANOVA, Mann-Whitney U tests, or Monte Carlo permutation testing, as appropriate and indicated in the figure legends. In the text, figures, and figure legends, data are presented as mean ± SEM. In the figures, statistical significance is denoted by asterisks (*p < 0.05, **p < 0.01, ***p < 0.001).

## Data availability

NGS data that support the findings of this study are deposited in the Sequence Read Archive (https://www.ncbi.nlm.nih.gov/sra) under the accession number PRJNA1223949. The output of normZ analyses of gene target enrichment and depletion are provided in Supplementary Tables 1 and 2. Source data are provided with this paper.

## Code availability

The code written for this study is shared in an online repository (https://github.com/kerschensteinerd/shen_2025).

## Acknowledgments

We thank the patients and organ donors who participated in this study, as well as Dr. Gary Marklin, Delaney Alyse, Brice Love, Abbi Van Camp, and Erica Hinterser at Mid-America Transplant for providing access to donor posterior poles. We are also grateful to Drs. Philip Custer, Kisha Piggott, and Josh Morgan from the Department of Ophthalmology and Visual Sciences at Washington University for providing retinas from enucleation surgeries for choroidal malignant melanoma. We thank Dr. Jenna Krizan from the Bright Center for Human Vision at Washington University for coordinating the procurement of human tissue, and Dr. Florentina Soto for producing the *AAV-PhP.eB-CAG-GFP* virus. We appreciate the expert technical assistance of Mike Casey, Ssu-Yu Chou, and Elizabeth Byrne. This work was supported by funding from the Hope Center for Neurological Disorders (P.A.R. and D.K.), that National Institutes of Health (EY027411, EY034001, and EY026978 to D.K., EY036368 to P.A.R, and EY002687 to the Department of Ophthalmology and Visual Sciences) and Research to Prevent Blindness (Career Development Award to P.A.R and unrestricted funds to the Department of Ophthalmology and Visual Sciences). This work was supported by the Hope Center Viral Vectors Core at Washington University School of Medicine.

## Author contributions

N.S., P.A.R., and D.K. conceived this work and designed the experiments. N.S., E.G.H, S.H, P.A.R., and D.K. conducted and analyzed the genome-wide CRISPR screen. M.F. measured and analyzed the pupillary light responses. N.S. conducted and analyzed all other experiments. N.S., P.A.R., and D.K. wrote the manuscript with input from all co-authors.

---

**Extended Data Fig. 1.**
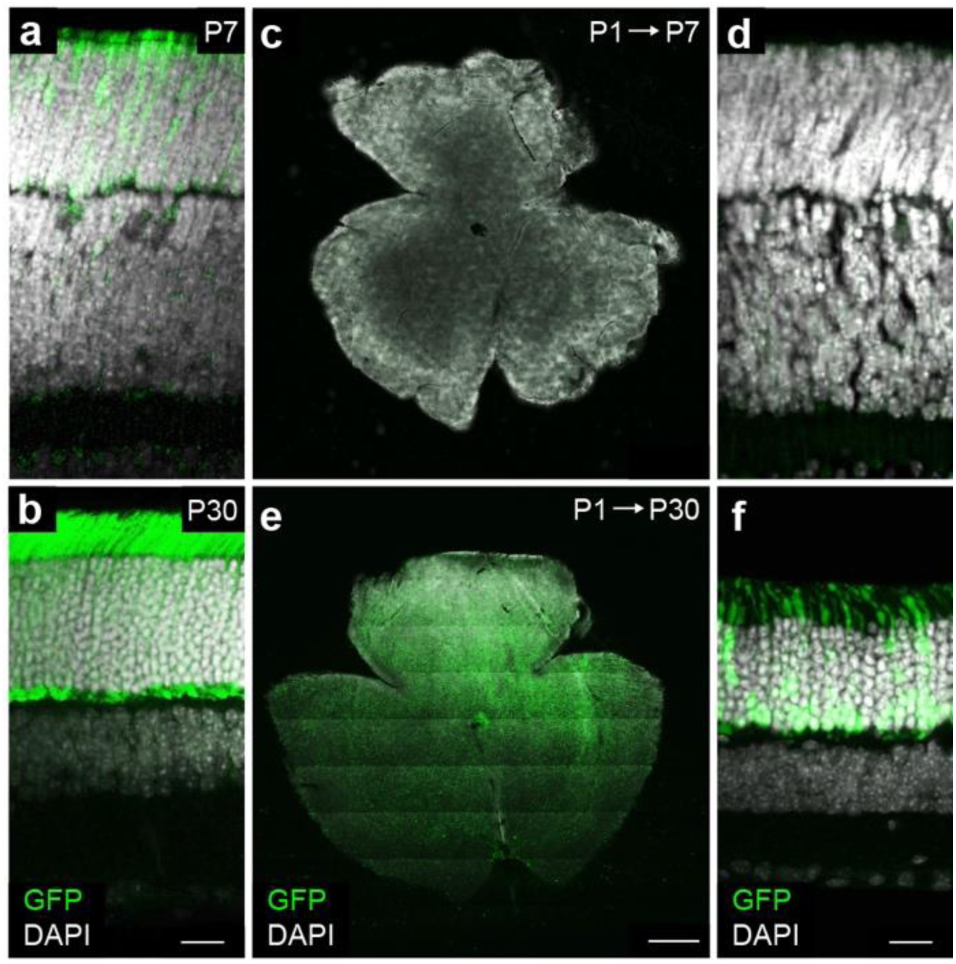
Time course of Cas9 and LV expression. **a,b,** Representative images of vertical slices from retinas of *Nrl-Cas9* mice at P7 (**a**) and P30 (**b**) show increasing Cas9 expression visualized via a GFP-tag included in the transgenic allele ^126^. Scale bar, 20 µm. **c-f,** Retinal whole mounts (**c,e**) and vertical slices (**d,f**) from mice injected subretinally with *LV-EF1α-GFP* at P1 and analyzed at P7 (**c,d**) or P30 (**e,f**). Scale bar, 500 µm in (**c,e**) and 20 µm in (**d,f**).

**Extended Data Fig. 2.**
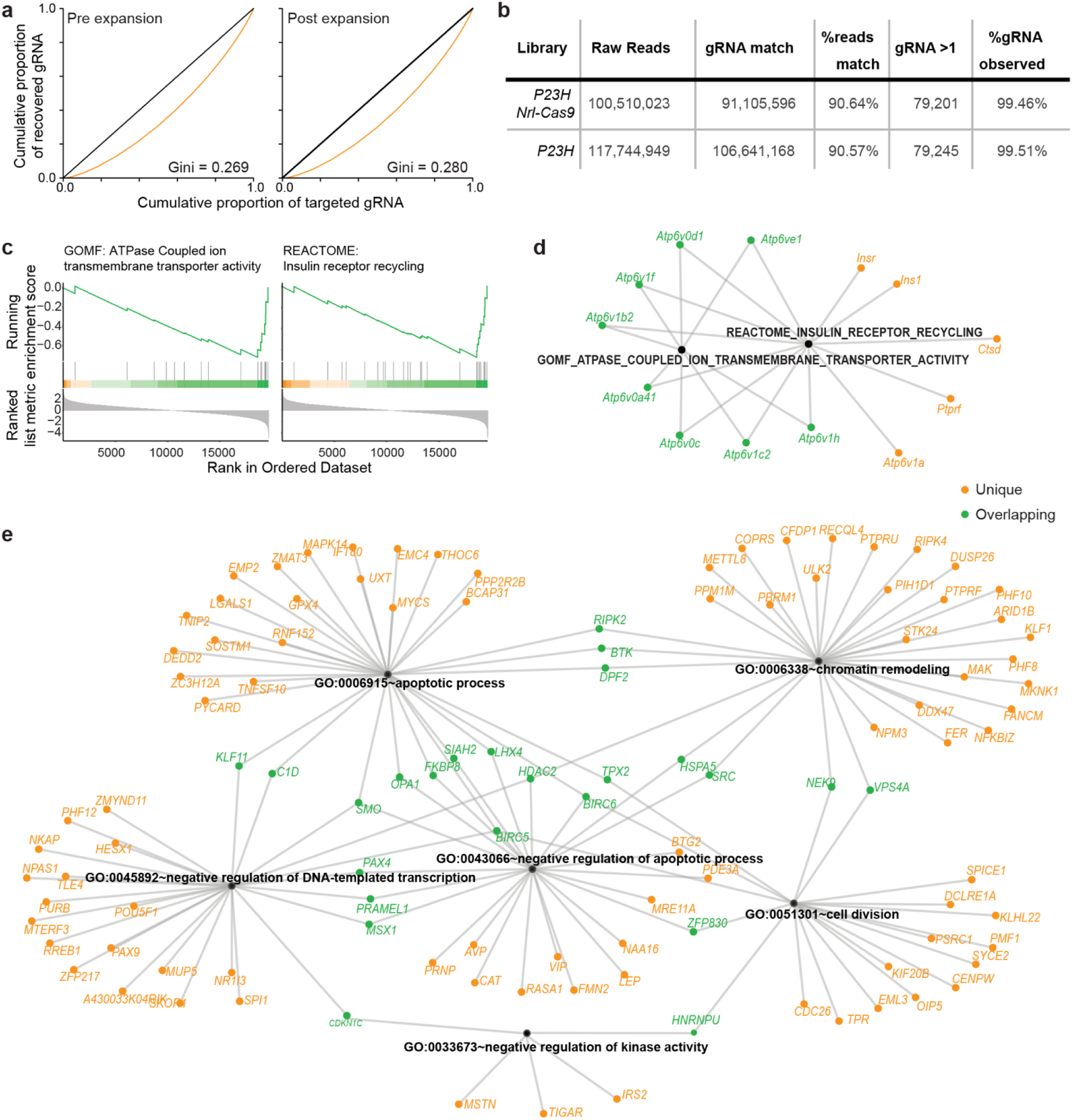
CRISPR screen reagent validation and outcome analysis. **a,** sgRNA library showed even coverage of sgRNA after expansion as measured by Gini coefficient. **b,** Coverage statistics of primary screen sequencing data. **c–e,** Enrichment analyses identifying enrichment of specific cellular pathways enriched in depleted gene set. GSEA (**c,d**) highlighted two overlapping functional categories while DAVID analysis (**e**) of genes normZ < -2 identified five largely distinct cellular functions.

**Extended Data Fig. 3.**
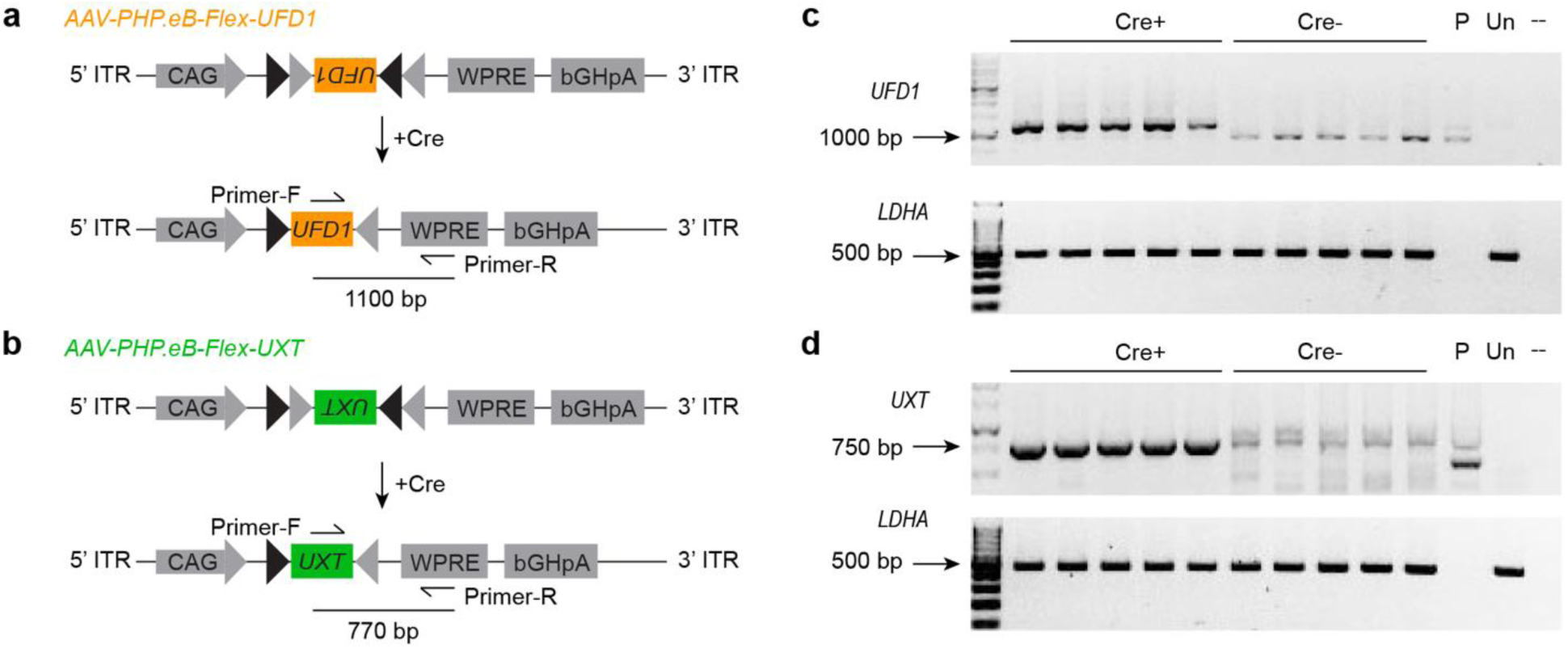
RT-PCR analysis of Cre-dependent gene augmentation. **a,b,** Schematics of the AAV constructs and primers designed to detect Cre-dependent overexpression of *UFD1* (**a**) and *UXT* (**b**). **c,d,** Reverse transcription-polymerase chain reaction (RT-PCR) validation of *UFD1* (c) and *UXT* (d) overexpression in AAV-injected *P23H Nrl-Cre* retinas. P = *p-AAV-CAG-Flex-UFD1/UXT* plasmid; Un: retinas without AAV injection.

**Extended Data Fig. 4.**
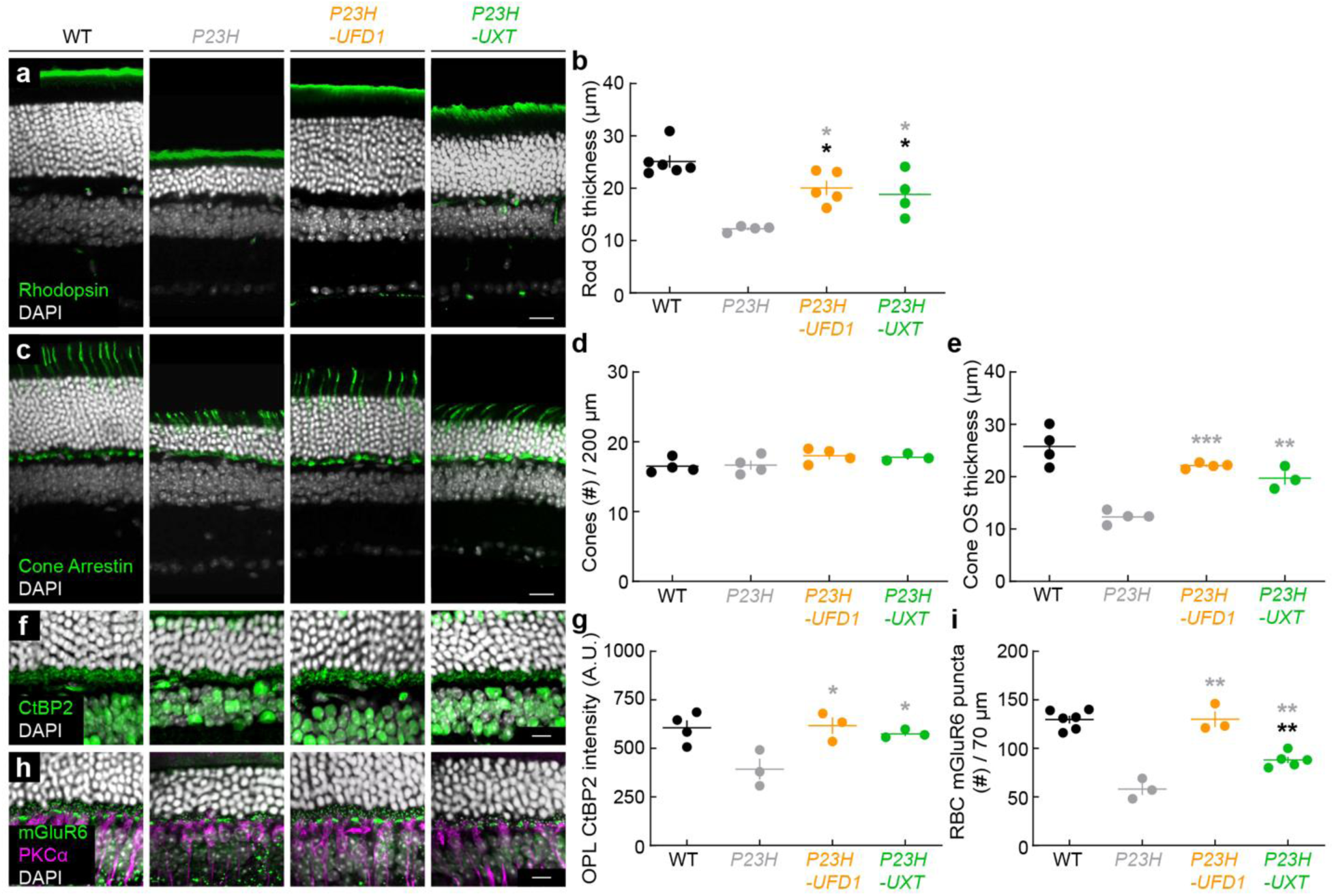
*UFD1* and *UXT* augmentation improve photoreceptor survival, morphology, and connectivity in the dorsal *P23H* retina. **a,** Representative confocal images of dorsal retinal sections from wild-type (WT), untreated *P23H*, or *P23H* treated with *UFD1* (*P23H-UFD1*) or *UXT* (*P23H-UXT*), stained for rhodopsin (green) and DAPI (white). Scale bar, 20 µm. **b,** Quantification of rod outer segment (OS) thickness. **c,** Representative images of dorsal retinal sections from WT, *P23H*, *P23H-UFD1*, and *P23H-UXT* stained for cone arrestin (green) and DAPI (white). Scale bar, 20 µm. **d,e,** Summary data for cone density (**d**) and OS thickness (e). **f,** Representative images of dorsal retinas stained for CtBP2 (green) and DAPI (white), labeling presynaptic ribbons. Scale bar, 10 µm. **g,** Quantification of CtBP2 immunofluorescence in the outer plexiform layer (OPL). **h,** Representative images of dorsal retinas stained for mGluR6 (green), PKCα (magenta), and DAPI (white). Scale bar, 10 µm. **i,** Quantification of mGluR6 puncta at rod bipolar cell dendrites. For (**b,d,e,g,i**), absence of an asterisk indicates p ≥ 0.05; black asterisks (*p < 0.05, **p < 0.01) indicate significance vs. WT, while gray asterisks (*p < 0.05, **p < 0.001, ***p < 0.001) indicate significance vs. *P23H* by one-way ANOVA.

